# Ligand Induced Conformational and Dynamical Changes in a GT-B Glycosyltransferase: Molecular Dynamic Simulations of Heptosyltransferase I Apo, Binary and Ternary Complexes

**DOI:** 10.1101/2021.06.16.448588

**Authors:** Bakar A. Hassan, Jozafina Milicaj, Yuk Y. Sham, Erika A. Taylor

**Affiliations:** Department of Chemistry, Wesleyan University, Middletown, CT 06459, USA; Bioinformatics and Computational Biology Program, University of Minnesota, Minneapolis, MN 55455, USA; Department of Integrative Biology and Physiology, Medical School, University of Minnesota, Minneapolis, MN 55455, USA

## Abstract

Understanding the dynamical motions and ligand recognition motifs of specific glycosyltransferase enzymes, like Heptosyltransferase I (HepI), is critical to discerning the behavior of other carbohydrate binding enzymes. Prior studies in our lab demonstrated that glycosyltransferases in the GT-B structural class, which are characterized by their connection of two Rossman-like domains by a linker region, have conservation of both structure and dynamical motions, despite low sequence conservation, therefore making discoveries found in HepI transferable to other GT-B enzymes. Through a series of 100 nanosecond Molecular Dynamics simulations of HepI in apo enzyme state, and also in the binary and ternary complexes with the native substrates/products. Ligand free energy analysis allowed determination of an anticipated enzymatic path for ligand binding and release. Principle component, dynamic cross correlation and network analyses of the simulation trajectories revealed that there are not only correlated motions between the N- and C-termini, but also that residues within the N-terminal domain communicate via a path that includes substrate proximal residues of the C-terminal domain. Analysis of structural changes, energetics of substrate/products binding and changes in pK_a_ have elucidated a variety of inter- and intradomain interactions that are critical for catalysis. These data corroborate and allow visualization of previous experimental observations of protein conformational changes of HepI. This study has provided valuable insights into the regions involved in HepI conformational rearrangement upon ligand binding, and are likely to enhance efforts to develop new dynamics disrupting enzyme inhibitors for GT-B structural enzymes in the future.

## Introduction

Enzymes that are involved in the transfer of sugar moieties, including glycoside hydrolases, glycosyltransferases, polysaccharide lyases and glycan phosphorylases, are critically important for specific functions such as bacterial biofilm formation, SARS-CoV-2 host recognition, regulation of cell cycle and tumor initiation.^[1–5]^ Due to the importance of glycosylation for processes ranging from energy storage to the biosynthesis of natural product therapeutic agents, significant advances have been made in the investigation of glycosyltransferase (GT) enzymes; however, additional research on this important class of enzymes is necessary for enhancing efforts to discover inhibitors and to employ these enzymes for other commercial synthesis applications.^[6, 7]^

GTs have been classified into 111 different enzyme families within the Carbohydrate-Active enZYmes Database^[8]^ (CAZY; http://www.cazy.org/GlycosylTransferases.html;), with families being defined utilizing sequence, structure and molecular mechanism. Within the database of over 120,000 proteins, structural information exists for 288 proteins, which allows these families to be classified into structural classes GT-A, GT-B and GT-C (representing 31, 28 and 11 families, respectively);^[9]^ though there are some GT families with unknown structures (38 families), others adopt previously identified folds (I.e. Family 51 adopts the lysozyme fold), while others have been classified as GT-D and GT-E (family 101 and 26, respectively). Various researchers have contributed to our understanding of the reaction mechanism of glycosyltransferase enzymes, and that all of the GT folds studied to date allow for the catalysis of sugar transfer with either retention or inversion of the stereo-configuration at the anomeric carbon.^[10, 11]^ While research to date supports a simple general acid-base catalyzed nucleophilic substitution mechanism for catalyzing the reactions with an overall inversion of stereochemistry, multiple reaction mechanisms have been observed for GTs that catalyze retention reactions. While some have proposed a double displacement mechanism to afford an overall retention of anomeric stereoconfiguration,^[12]^ only two enzymes of the GT-A scaffold have been shown to have a nucleophile present in the proper orientation to allow this. In the majority of retaining GT enzymes, including MshA^[13]^ (a GT-B enzyme) evidence supports the enzymes using a S_N_i mechanism without the involvement of an active site nucleophile.^[14–16]^

Only a small fraction of the GT families represented in CAZY have available crystal structures.^[10, 17]^ Furthermore, even fewer of these structures have either ligand or both ligands present. The difficulty of crystallizing these enzymes with their substrates has slowly been overcome with unique strategies including co-crystallization with fluorinated donors^[18–21]^ and functionally equivalent acceptor analogues.^[22]^

In recent decades, there have only been a handful of GTs that have been simulated. The sparse number of available crystal structures and the even lower number of liganded complexes has hindered the effort towards understanding these enzymes on an atomistic and dynamic scale. Both GT-As and GT-Bs have been simulated on relatively short timescales (100-250ns). Simulations of GT-Bs PglH,^[23]^ alMGS^[24]^ and GumK^[25]^ have provided insights into the ligand interactions and membrane related behavior that would otherwise be difficult to elucidate. Similar work has been performed with several GT-As.^[26, 27]^ In addition, simulations have been used in tandem with homology modeling to further study GTs when crystal structures are unavailable.^[28–31]^

Heptosyltransferase I (HepI) is Glycosyltransferase in the GT9 family with a characteristic GT-B fold. It consists of two domains with β/α/β Rossman-like folds connected by a linker (Figure 1A).^[18]^ HepI is involved in the lipopolysaccharide biosynthetic pathway and transfers a seven-carbon heptose sugar via ADP-L-glycero-β-D-manno-heptose (ADP-Hep) to the first 3-deoxy-D-*monno*-oct-2-ulosonic acid (Kdo) of the membrane anchored Kdo_2_-Lipid A (Figure 1B).^[32–35]^ The reaction produces heptosylated Kdo_2_-Lipid A (H-Kdo_2_-Lipid A) where the stereochemistry at the anomeric carbon of the heptose sugar donor is inverted, therefore, classifying HepI as an inverting GT-B. Like other GT-Bs the donor and acceptor are greater than 10 Å apart in the open conformation and these enzymes undergo a global conformational change that brings the reaction centers to within a tolerable range for catalysis. Several other GT-Bs have solved structures that show this global conformational change and these include GtfA, MshA and Glycogen synthase.^[36–38]^ It has been previously shown that HepI potentially undergoes a conformational change upon binding of the acceptor ligand, and this was determined with Tryptophan reporters.^[39, 40]^ In addition, pre-steady state kinetics has shown the rate limiting step to be something other than catalysis and this is believed to be the conformation change induced by acceptor binding.^[41]^

**Figure 1:**
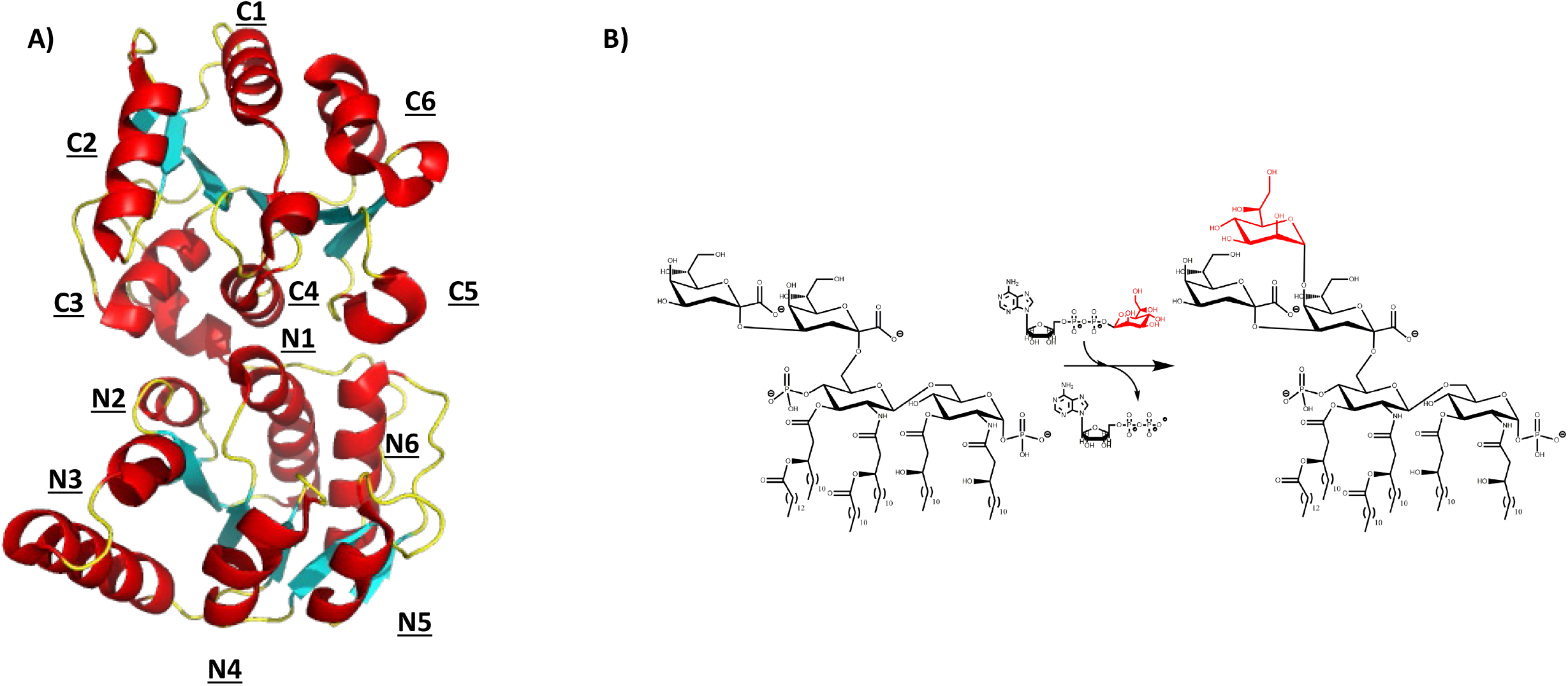
**A)** Structure of HepI (PDB: 2GT1) labeled by order of secondary structure in respective domains. **B)** Reaction catalyzed by Heptosyltransferase I.

Recently we have performed long timescale (microsecond) simulations of HepI and a distantly related GT-B from vancomycin antibiotic biosynthetic pathway, GtfA.^[42]^ While HepI has not been structurally characterized in the “closed” conformation, GtfA has been crystallized in the “closed” conformation with both ligands present. We showed that when the ligands are removed, GtfA returns to its unbound state but maintains dynamic modes that are important for “closed” conformation. The modes are also present in HepI in the unbound state, but to a lower degree, which suggest that these quasi-harmonic modes are conserved among GT-Bs and that there are dynamic changes upon ligand binding. Recently, a crystal structure of HepI with a mostly deacylated acceptor substrate and a non-hydrolyzable glycoside acceptor analoguewas solved.^[22]^ This combined with previous solved structures of HepI with a fluorinated donor analogue, has now provided a working model for the fully liganded ternary complex.^[18]^ Molecular dynamic simulations of HepI modeled with both ligands could provide insights into the dynamics and energetics of substrate binding to aid in the future design of inhibitors that could act as effective antimicrobials.

## Methods

### Multiple Sequence Alignment

Multiple sequence alignment for *E. coli K12* HepI was obtained through ConSurf as previously described.^[43, 44]^ Briefly, a reference sequence was obtained from the pseudo ternary complex of HepI (PDB:6DFE) and a protein BLAST with uniprot90 server yielded 2028 unique sequences that have 95-35% sequence identity to the reference. Sequences were filtered by E values with a cutoff of 0.0001 and 150 of those sequences were chosen by compiling a list with sequences of the every 15^th^ index to equally sample the whole list of homologues. Clustal Omega was used for multiple sequence alignment of 150 representative sequences and Maestro was used to construct a logo plot.^[45]^

### Modeling

All HepI structural models used in this study were utilizing previous crystal structures or a hybrid of multiple crystal structures (Supplemental table 1). The apo model was simulated utilizing a previously solved structure (PDB:2TG1).^[18]^ The fully ligated substrate model was constructed using the pseudo-ternary complex (PDB:6DFE)^[22]^ to provide the protein and sugar acceptor geometries and interactions, while the sugar donor carbamate analogue was replaced with the fluorinated sugar donor from a previously solved binary complex (PDB: 2H1H)^[18]^ and the fluorine on the sugar donor analogue at the C2 position was replaced by a hydroxyl group with the inversion of stereo-configuration to match that of the native substrate. The sugar acceptor was an analogue of Kdo_2_-Lipid A with the acyl chains removed from the O and N positions of the N-acetyl-Glucosamine. A singular acyl chain was kept at each one of the N positions of the N-acetyl-Glucosamine, therefore, this sugar acceptor will be referred to as fully-deacylated Kdo_2_-Lipid A (FDLA). Products were modeled based on the fully ligated substrate model, above, with the transfer of the heptose moiety from ADP to the FDLA to reflect their conversion to the products, to form adenosine disphosphate (ADP) and fully-deacylated heptosylated-Kdo_2_-Lipid A (FDHLA). The binary complex of the sugar donor (ADP-Hep) and the product (ADP) were modeled with previously solved structures PDB:2H1H and PDB:2H1F, respectively.^[18]^ The binary complex of the sugar acceptor (FDLA) was modeled with the previously solved structure PDB: 6DFE^[22]^ with the removal of the sugar donor carbamate analogue. Similarly, the binary complex of the sugar acceptor product (FDHLA) was modeled with the previously solved structure PDB: 6DFE^[22]^ with the modification of the FDLA as described above and subsequent removal of the sugar donor carbamate analogue.

### Molecular Dynamic

All molecular dynamic simulations were performed with GROMACS-2020.2^[46, 47]^ and the Amber99SB^[48]^ forcefield. Ionization states for titratable sidechains were determined with PROPKA3.^[49, 50]^ All systems were solvated with a transferrable intermolecular potential with 3 points (TIP3P)^[51]^ explicit solvent model in a cubic box with a 10 Å buffer region and electroneutralized with 0.150 M NaCl counterions. Equilibration was performed with harmonic restraints (1000 kJ/mol/nm^2^) on heavy atoms with a stepdown equilibration that involves removal of restraints from sidechains and then backbone over the course of 10 ns. Energy minimization was performed with the steepest descent algorithm. The system was equilibrated for 1 ns under isochoric/isothermal conditions (NVT) and a subsequent 1 ns equilibration under isobaric/isothermal conditions (NPT). Temperature and pressure were regulated with the Berendson thermostat/barostat.^[52]^ Production simulations were carried out at 300 K and 1 atm (NPT ensemble) for 100 ns with a time step of 2 fs in triplicate. Temperature and pressure were maintained via v-rescale and Berendson coupling respectively.^[52, 53]^ Short range nonbonded interactions were calculated with a cutoff of 1.0 nm and long-range electrostatic interactions were calculated with particle-mesh-Ewald with a fourth order cubic interpolation and 1.6 Å grid spacing. Bonds were constrained with the LINCS^[54]^ method. Ligand charges and atom types were assigned with the AM1-BCC model and the second generation Generalized Amber Forcefield (GAFF2), respectively.^[55, 56]^ This was accomplished via ANTECHAMBER from AMBERTOOLS20 package and the ligand files were subsequently converted to GROMACS compatible file format with the ACPYPE tool.^[57, 58]^ Simulations of the binary and ternary complexes with donor product (ADP) were unstable with ADP reproducibly leaving the active site within 20 nanoseconds of the start of the simulation, therefore, a 2.5 Å distance restraint between the hydrogens of the primary amine at 6 position of the adenosine ring on the ADP and backbone carbonyl oxygen of Met242 was used to keep the ADP in place. This Hydrogen bonding interaction occurs in all the structures with donor substrate (ADPHep) or donor products present (ADP) in the active site and was believed to be the best way to keep the product in the active site without restricting its conformational flexibility.

### RMSD, RMSF, PCA and Network Analysis

Ligand interaction diagrams were made in Maestro. Minimum distances between protein residues and ligands over the course of 100 ns were calculated in GROMACS. Root mean square deviations (RMSD) and root mean square fluctuations (RMSF) of the backbone and C_α_ atoms respectively, were calculated in GROMACS over the course of 100 ns and averaged from three separate simulations. RMSD and RMSF plots were subsequently generated in Python. ΔRMSF was determined for each condition with respect to the averaged apo RMSF and shaded regions correspond to the average standard deviation difference. Principal component analysis (PCA) and network analysis were both performed in R with the bio3D package on one representative trajectory and have been extensively described elsewhere^[59–61]^. Briefly PCA involves the diagonalization of the covariance matrix into their component eigenvalues and eigenvectors. The eigenvectors represent the set of possible structures (principal components) and the eigenvalues represent the covariance of those structures. This reduces the dimensionality of the data set to explore the conformers of the trajectory. The “motion” associated with each of the principle component can be further explored by mapping the extreme points of the principal component on the average structure and interpolating to generate a movie that describes that principal component. The coordinated dynamics between residues can be determined by calculating a correlation between the displacement of two atoms relative to their average position (Equation 5).^[62]^ A per residue dynamic cross correlation matrix (DCCM) can be determined by calculating this correlation value for C_α_ atoms relative to one another. This provides a metric in which two atoms can be described as having motions in an identical direction (i.e. positively correlated) or opposite direction (i.e. negatively correlated).

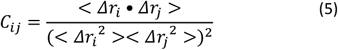

To determine shortest path between residues, each residue was considered a node. Every node was connected to another node via edge if they are in contact. The edges are weighted by their dynamic cross correlation value. The path length is the sum of the weights and the shortest path is determined by finding the smallest path length between residues of interest.^[61]^ A correlation cutoff of 0.5 was used for constructing the DCCM. In addition, PCA and network analysis, ligands were ignored and only Cα were considered. All molecular graphic models were made in PyMol.

### Energetics

The molecular mechanics Poisson Boltzmann solvent accessibility (MMPBSA)^[63]^ end point method for estimating the binding free energy of ligand to a macromolecule has been extensively described elsewhere.^[64–66]^ Briefly, the binding free energy of a receptor ligand complex is the free energy difference of the complex from the receptor and ligand. The free energy of each of these components is the sum of the bonded and nonbonded energy of the system. In addition, the free energy of the solvation is estimated as the sum of the polar contribution derived with the Poisson Boltzmann implicit solvent model and the nonpolar contribution is calculated with the solvent accessible surface area. The entropic contribution is estimated via normal mode analysis.

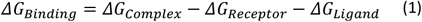

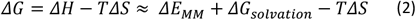

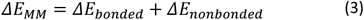

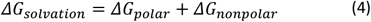

The MMPBSA method was implemented in AMBERTOOLS 20 with the MMPBSA.py script^[67]^ The fully ligated substrate/product complex trajectory was used for to determine the energetics of each step. From the substrate/product complex trajectory, 20 representative frames were chosen and an internal dielectric of 4 best described the charge distribution of HepI. Each free energy was calculated in triplicate and reported with a standard deviation.

## Results

### Differential Dynamic Flexibilities in the Apo and Liganded Complexes of HepI

Simulations were performed of Heptosyltransferase I on the 100 nanosecond timescale as the apo enzyme, and the (1) ADP-Hep•HepI, (2) FDLA•HepI, (3) ADP-Hep•FDLA•HepI, (4) ADP•H-FDLA•HepI, (5) ADP•HepI, (6) H-FDLA•HepI (7) ADP-Hep•FDLA-H•HepI (D13+H), and (8) ADP•FDHLA•HepI (D13+H) complexes. To evaluate the overall stability of our systems, we monitored the average backbone root mean square deviation (RMSD) and the average radius of gyration (R_gyr_). The backbone root mean square deviation (RMSD) of the apo, substrate and product ternary complexes are 1.70 ± 0.25 Å, 1.75 ± 0.31 Å, and 1.85 ± 0.31 Å respectively (Supplemental Table 2). The binary complex RMSDs are within 0.2 Å of their ternary complex counterpart. The R_gyr_ for the apo, substrate and product complex are 21.11 ± 0.16 Å, 21.21 ± 0.12 Å, and 21.36 ± 0.23 Å respectively. Similarly, the binary complexes are within approximately 0.4 Å of their ternary counterpart. The RMSD and R_gyr_ quickly stabilize and no significantly observable deviations occur after the first 10 nanoseconds (Figure 2A; Supplemental Figure 1). The apo simulation reveals three regions that exhibit Cα root mean square fluctuations (RMSF) that are greater than 1.5 Å (not including the dynamic tail; Figure 2B, Supplemental Figure 2; Supplemental Table 2). These regions include residues in the N3 (61-68), C2 (216-219) and C6 (316-320). The substrate ternary complex has residues with RMSF values greater than 1.5 Å in the N3 (62-67), C1 (188-189), C5 (283-284) and C6 (299,317-320). The ternary product complex has residues with RMSF values greater than 1.5 Å in the N3 (61-68), N4 (103), N6 (135,156), C1 (188-189,206), C2 (218), C5 (280-281, 287-291) and C6 (298-318). The simulations of the binary complexes have fluctuations in similar regions as their ternary counterparts, where the substrate complexes are generally less dynamic than the product complexes. The absolute per residue fluctuations, as provided by the RMSF, can mask small changes or regional changes in relative fluctuations. ΔRMSF provides a better insight into the relative and regional changes among our simulations. Similarly to the RMSF, in the ΔRMSF we see the greatest standard deviation of relative motion in the N3, N4, and N5 (Figure 2C; Supplemental Figure 3). These high standard deviations most notably occur in the ADP•FDHLA•HepI product ternary complex with the protonated Asp-13.

**Figure 2:**
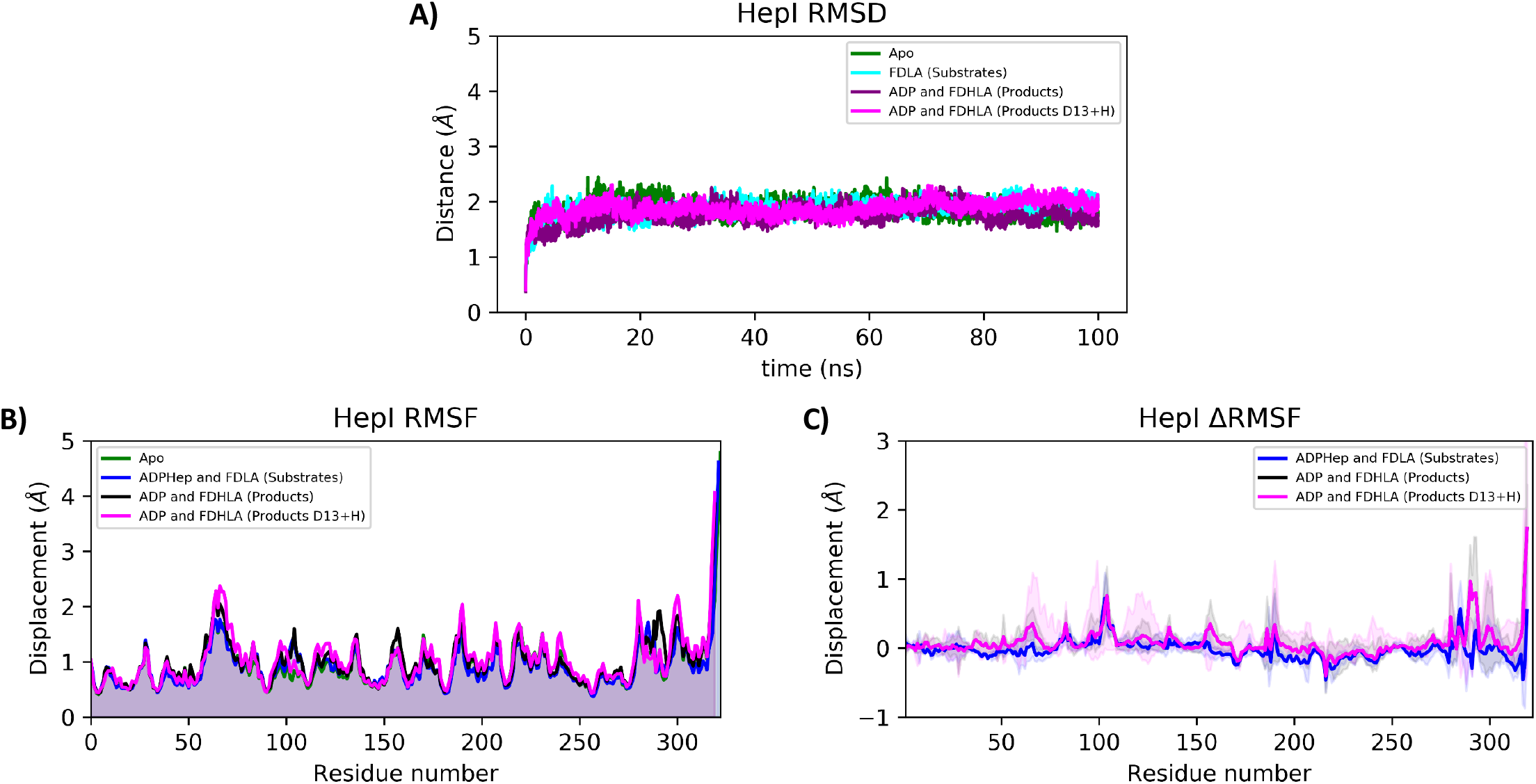
(**A**) Backbone RMSD, (**B**) C_α_ RMSF and (**C**) C_α_ ΔRMSF of HepI Apo (green), substrate (blue) and product ternary complexes (black for the product complex with Asp13 deprotonated and magenta for the product complex with Asp13 protonated). For C_α_ ΔRMSF solid lines are average differences relative to apo (i.e. RMSF_substrates_-RMSF_apo_) and shaded region is the standard deviation of the average difference. Positive values indicate those residues are more flexible relative to HepI apo.

### Protein-Ligand Interactions and Binding Affinities of HepI Complexes

HepI has two domains, with the N-terminal domain that binds the acceptor (FDLA) and the C-terminal domain that binds the donor (ADP-Hep). Previously solved structures^[18, 22]^ have identified a hydrophobic pocket adjacent to the adenosine ring in the C-terminal binding pocket. Furthermore, it has been suggested that a hydrogen bond between the N6 of the adenosine ring and the backbone carbonyl of Met242 facilitates substrate capture along with hydrogen bonds between the ribose hydroxyls and Glu222. The pyrophosphate is stabilized through electrostatic interactions with basic residues like Lys192 and hydrogen bonding with residues like Thr187-Thr188. The sugar acceptor binding site has been shown to be stabilized by a collection of electrostatic interactions with basic residues in the N3 and N5 loops. Furthermore, through mutagenesis, Asp13 was implicated as the catalytic base and is strictly conserved across all sequenced homologues. This residue is in the N terminal domain and is located adjacent to the C5 hydroxyl of FDLA to facilitate a proton transfer for catalysis.

Examination of the ligand complexes revealed constellations of additional residues involved in ligand binding during the course of the simulations. In both the HepI binary and ternary complex simulations, the oxygens of the β-phosphate of ADPHep hydrogen bond with the backbone amides of residues Met11 and Gly12 in the N-terminal domain (Figure 3B & 3D). Additionally, the C-terminal domain backbone of Thr188 hydrogen bonds with the α-phosphate oxygens of ADPHep in both the binary and ternary complex (Figure 3B & 3D and Supplemental Figure 4A-B & 4E-F). The primary amine at 6 position of the adenosine ring on the ADPHep hydrogen bonds with the backbone oxygen of Met242 in these complexes. Lastly, the Heptose hydroxyl groups hydrogen bond with Lys192, Asp261, and His266 in both complexes. In the HepIoADP complex, in addition to the hydrogen bonding interactions with Met11, Gly12, and Met242, analogous to those described above for ADPHep, the α phosphate oxygens hydrogen bond with backbone amines of residues Gly263, Thr262 (in both the binary and ternary complexes)(Supplemental Figure 4I-J & 4M-O). The binary complex also forms a salt bridge between the sidechain of Lys192 and the α phosphate, while the sidechain of Arg60 also forms a hydrogen bond with the hydroxyl of the ribose; however in the ternary complex the interactions between Arg60 and ADP-Hep is missing, and Arg60 now interacts with the phosphate of FDHLA (Supplemental Figure 4L, 4Q).

**Figure 3:**
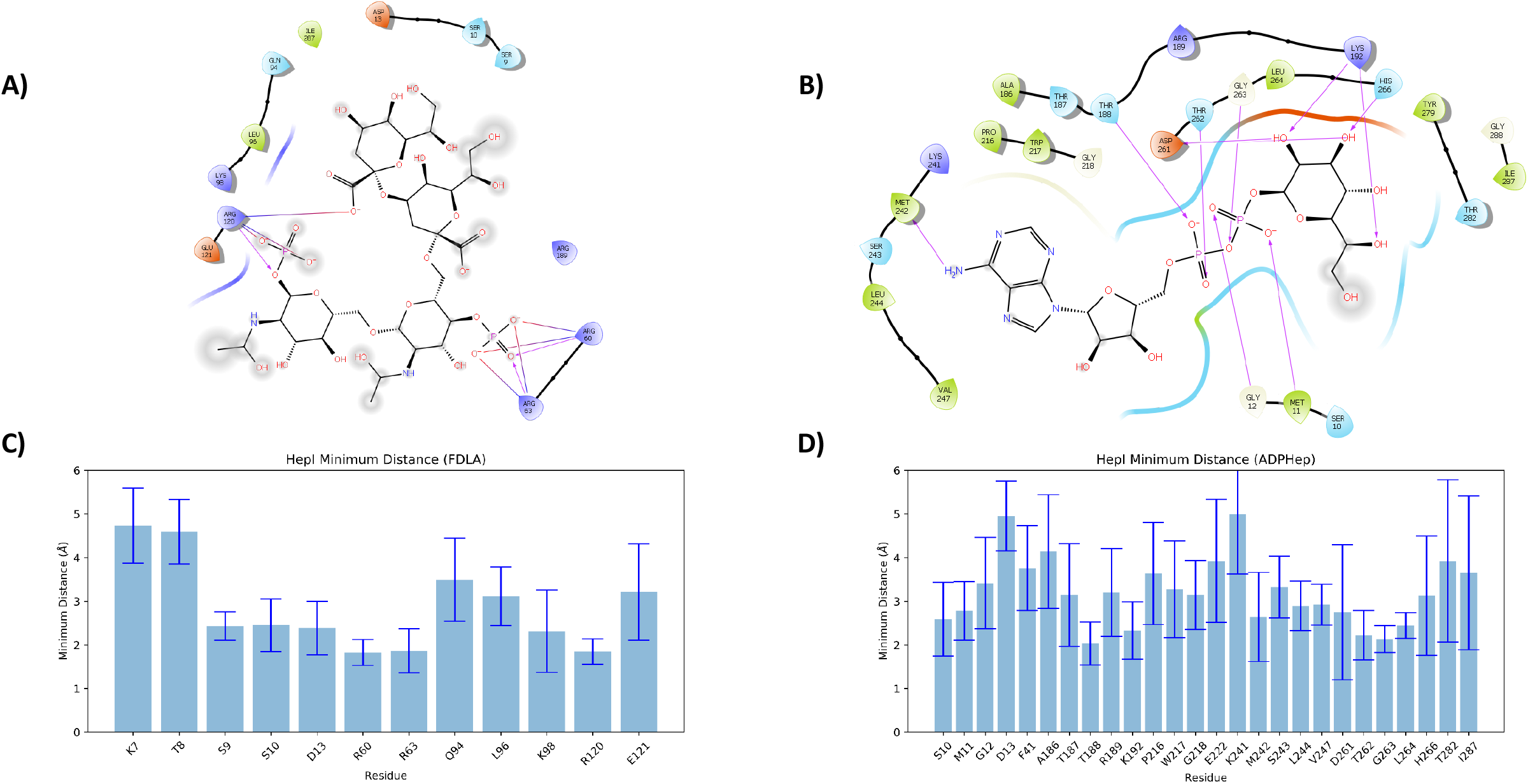
Ligand interaction diagram (**A**) FDLA and (**B**) ADP-Hep from ADP-Hep•FDLA•HepI (Substrates) ternary complex simulation. Bar plots of residues with average minimum distances of less than 5 Å from FDLA (**C**) and ADPHep (**D**).

As observed in ADPHep and ADP, FDLA maintains numerous interactions in both the binary and ternary complex simulations, including forming salt bridges between the FDLA phosphates with arginines and lysine in the N3- and N5-loops; specifically Arg60, Arg63, Lys98, Lys120 and Arg189 interact in both binary and ternary complexes (Figure 3A, 3C; Supplemental Figure 4C-D, 4G-H). FDHLA demonstrates similar interactions in both binary and ternary complexes as observed with FDLA, with the addition of a hydrogen bond between the sidechain of the Asp13 and the carboxylate of the second Kdo in the ternary complex and the C3 hydroxyl of the transferred heptose. Interestingly, ADP does not stay bound to HepI ineither the binary or ternary complexes, with it leaving the active site in the first quarter of the simulations without being constrained in the active site by a hydrogen bond to Met242, described in the methods. Simulations with the ADP-Met242 hydrogen bond, showed the ligand adopting poses consistent with those previously observed via protein crystallography, and are therevfore anticipated to be physiologically relevant. Using an MM-PBSA method, the binding free energy was calculated in AMBER for the association of each ligand to the protein/protein•ligand complex to generate a theoretical thermodynamic cycle (Figure 4). The estimated binding free energy of the ADPHep to HepI in the binary complex is −16.63 ± 3.19 kcal/mol and FDLA is −13.27 ± 3.15 kcal/mol. The binding free energy of FDLA to ADP-Hep•HepI is −6.93 ± 3.06 kcal/mol and ADP-Hep to FDLA•HepI is −10.29 ± 3.03 kcal/mol.

**Figure 4:**
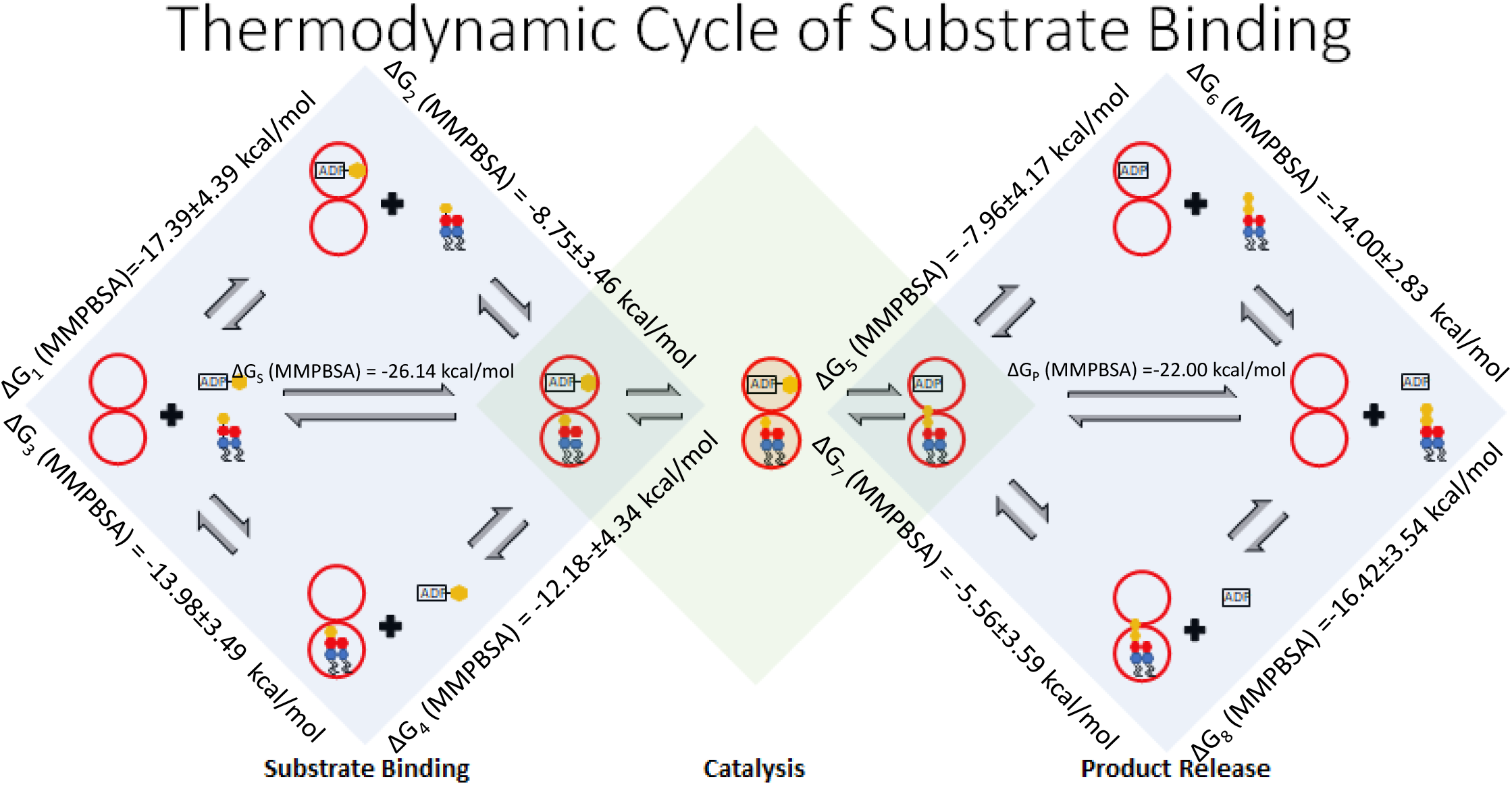
Thermodynamic cycle of substrate/product binding with binding free energies determined by MMPBSA for ADP-Hep to HepI (ΔG_1_), FDLA to ADP-Hep•HepI (ΔG_2_), FDLA to HepI (ΔG_3_), ADP-Hep to FDLA•HepI (ΔG_4_), FDHLA to ADP•HepI (ΔG_5_), ADP to HepI (ΔG_6_), ADP to FDHLA•HepI (ΔG_7_) and FDHLA to HepI (ΔG_8_).

### Local, Global and Correlated Conformational Motions of HepI Complexes

Ligand binding causes both local and global changes in HepI. One local change induced by the presence of substrates/products is the alteration of the ionization states of several sidechains, based upon the analysis of pKas using PROPKA3. Most importantly, the putative catalytic residue Asp13 has a pK_a_ of 4.46 in the apo enzyme, but shifts to 6.79 in the presence of the substrates (Supplemental Table 3). In the presence of the products, the pK_a_ of D13 shifts even further to 10.38. This highly purtubred pK_a_ is maintained in the binary complexes with either N-terminal ligand (FDLA or FDHLA) present. Lys7, which is located within hydrogen bonding distance of Asp 13,also exhibits a pK_a_ shift from 9.27 in the apo enzyme to 5.77 in the product ternary complex; this shift is not seen in the substrate complex.

Each of the simulation trajectories were analyzed to determine the global and local motions through principal component and dynamic cross correlation analyses to determine the impact of ligand(s) on the motions of HepI (Figures 5 and 6; Supplemental Table 4; Supplemental Figures 7 and 8). The total structural variance of HepI with the first three principle components ranged from 39-54.7% (Supplemental Table 4). To better understand these principle components, the extreme points were mapped onto the average structure and interpolated to generate a dynamic representation of each principle component. For the apo, the first three principle components have a near evenly dispersed variance of 19.2%, 13.9% and 11.1%. Whereas, for the substrate ternary complex, the first principle component predominates with 30.2% and the other two only accounting for 11.5% and 5.7%. The product ternary complex has a similar skew towards the first principle component with 29.1% and the other two contributing 14.1% and 10.7%, respectively. This difference is most evident with the product ternary complex where Asp 13 is protonated which has 40.7% variance for the first principle component and 11.6%, 5% for the second and third, respectively. The binary complexes follow similar distributions as their ternary complex counterpart.

**Figure 5:**
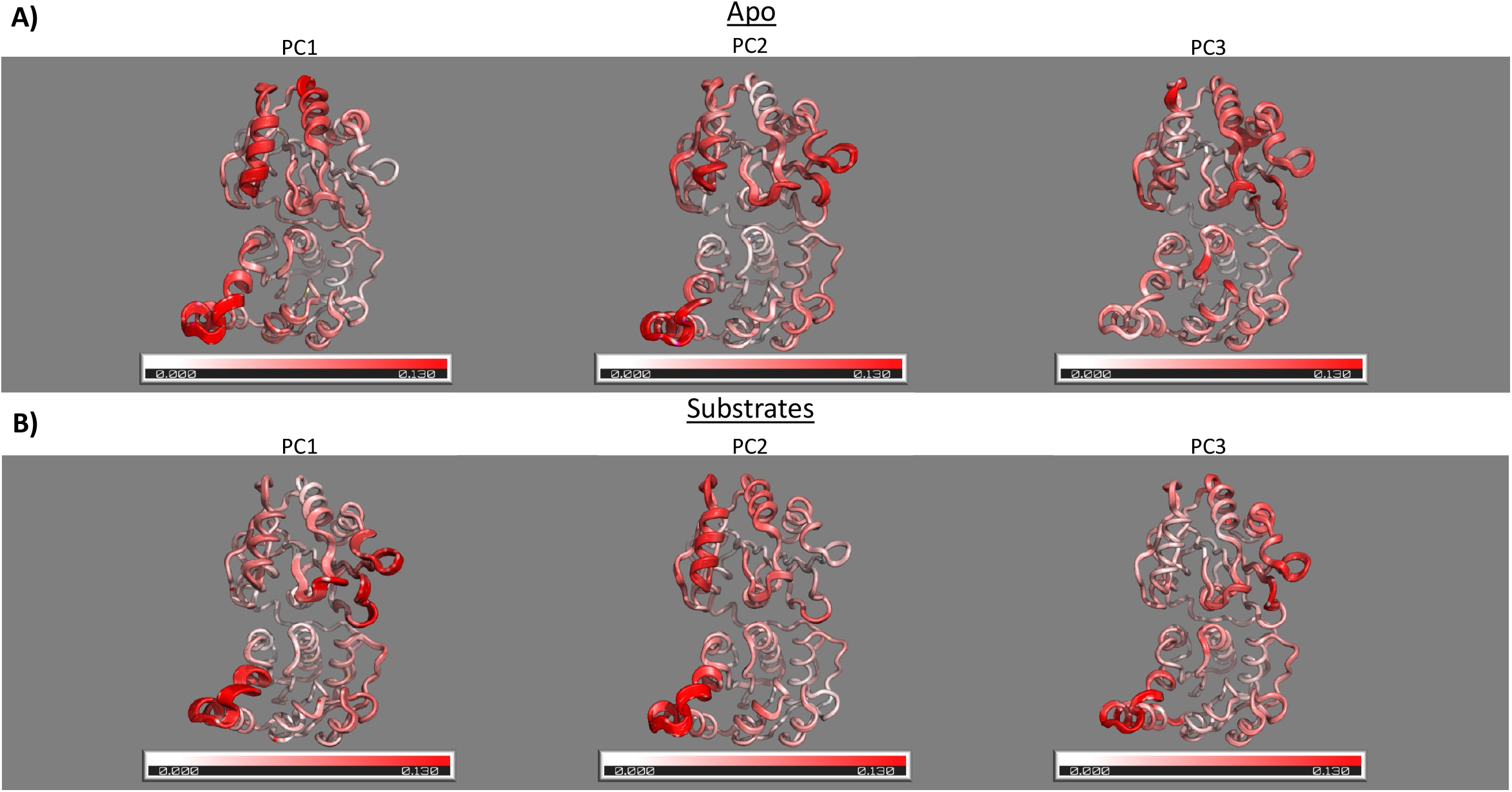
First three principle components of (**A**) HepI apo and (**B**) substrate complex. Interpolation of extremes points for each principle component onto the average structure gives rise to a motion that is represented by the thickness of the ribbon diagram. C_α_ root mean square fluctuation (RMSF) of each principle component is represented by the red color gradient with increasing color representing increasing motion.

Examination of the apo and ADPHep•FDLA•HepI complex DCCM simulations (Figures 6A-B and Supplemental Figures 8A, 8G, 8B, 8H) reveals an island of interdomain negatively correlated residues that correspond to coupled motion between residues in the N3 helix and the C2 helix. The substrate ternary complex DCCM illustrates that these interdomain negatively correlated motions between the N3 helix and C2 helix are enhanced by the presence of substrates and expanded to include negative correlations to the C1 helix. Additional islands of negatively correlated motions also appear between the N4/N5 helices and the C5/C6 helices, indicating that both sides of the two Rossman domains are engaged in anti-correlated motions. The product ternary complex simulation DCCM has a larger number of islands of positively correlated motions spread through both domains (Figure 6C and Supplemental Figure 8C, 8I); the coupled interdomain regions include the N1 helix and C2/C3 helix, N2 helix and C2/C3 helix, N3 helix and C3 helix, N4/N5 helix and C2 helix, N4/N5 helix and C4/C5 helix, while the coupled intradomain regions include N1 helix and N3 helix, N2 helix and N5 helix, C3/C4 helix and C5 helix. When Asp13 was protonated for the product ternary complex, there was a further enhancement of negatively correlated interdomain motions between N4 with C1/C4/C5/C6 and N5 with C1/C4/C5/C6 (Figure 6D). Neither the FDLA or FDHLA binary complexes display any islands of anti-correlated motions at a correlation cutoff of 0.5 (Supplemental Figures 8D-N). The ADP binary complex has modest negatively correlated regions between N5 helix and C4/C5 helix (Supplemental Figure 8J).

**Figure 6:**
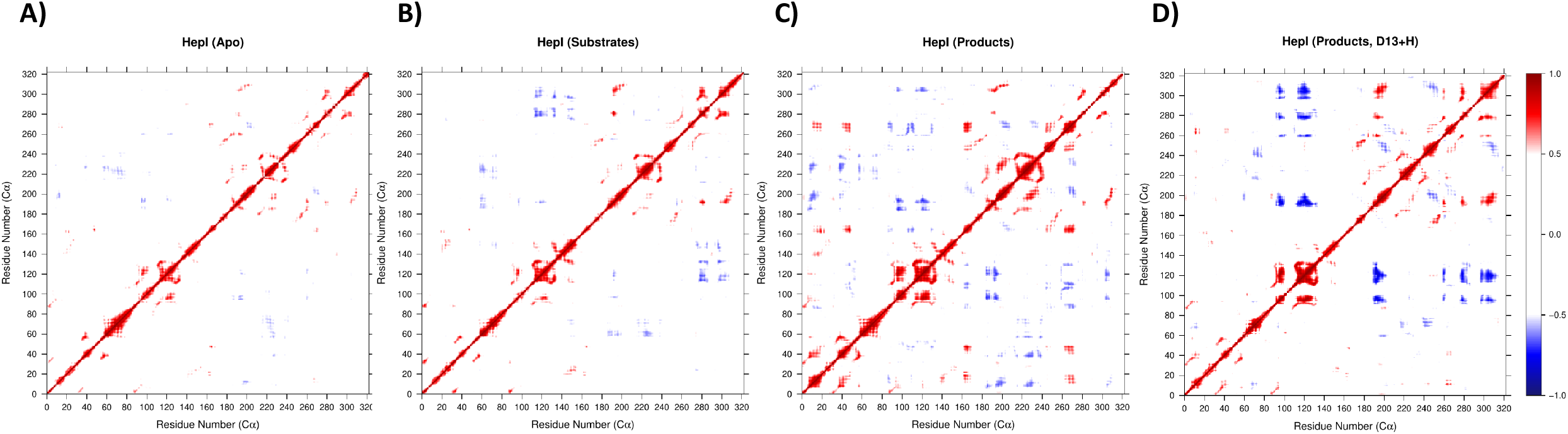
Dynamic cross correlation map of (**A**) HepI apo, (**B**) substrate ternary complex, (**C**) product complex with Asp13 deprotonated and (**D**) product complex with Asp13 protonated.

**Figure 7:**
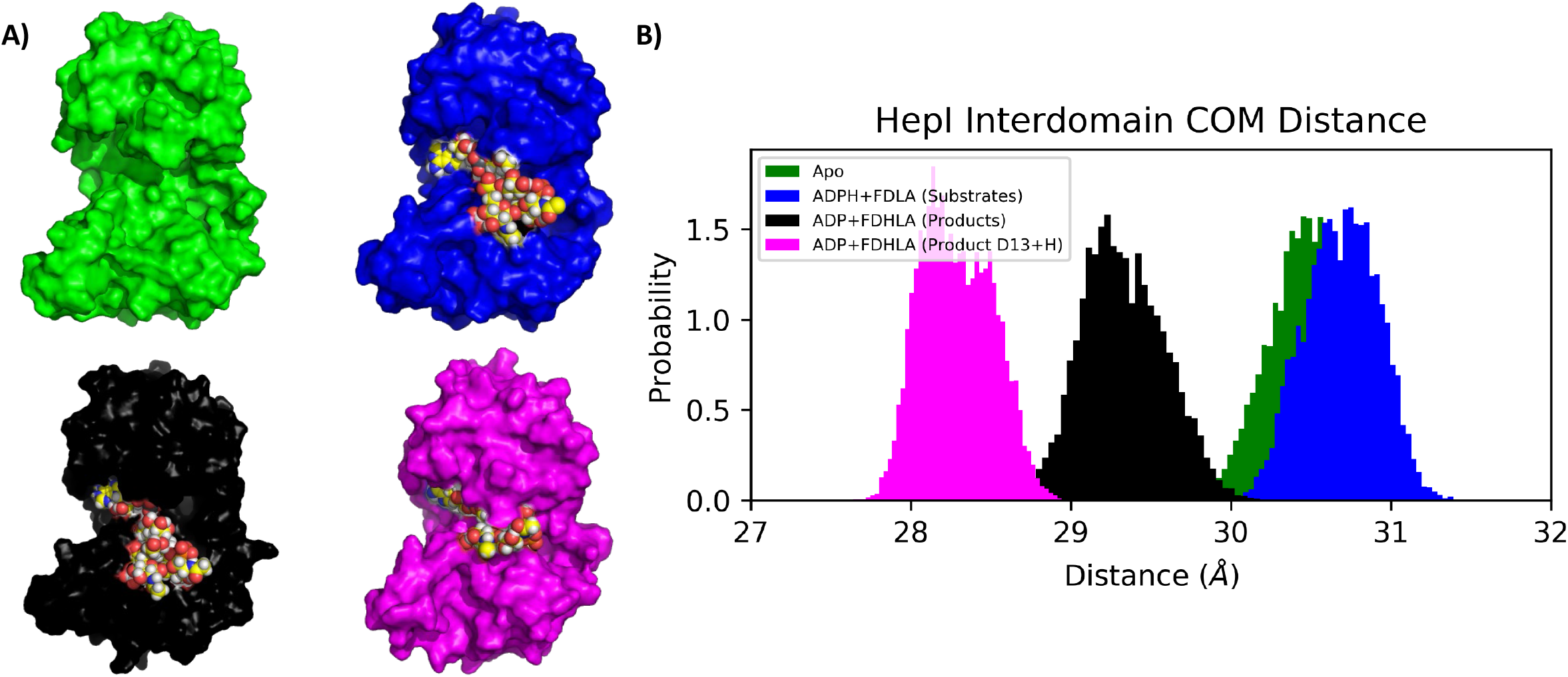
(**A**) Final frame surface representation of HepI simulations with space filling ligands and (**B**) probability distribution of center of mass distance between N and C termini of HepI in the apo (green), substrate (blue), product with deprotonated Asp13 (black) and product with protonated Asp13 complex simulations (magenta).

Dynamical network analysis allowed determination of the shortest paths of communication between residues that are involved in substrate binding or residues that are suspected to be involved in catalysis. These analyses reveal both intradomain and interdomain networks. In the apo state, Arg60 and Arg120 communicate through C2- and C1-alpha helical residues (Figure 8A; Supplemental Table 5; Supplemental Figure 9A). In the presence of substrates, Arg60 communicates through C1-helix residues and back down to Arg120 (Figure 8C; Supplemental Table 5; Supplemental Figure 9D). The shortest path for Asp13 (the catalytic base) to communicate with Met242 (a ligand of Adenosine in either ADPH or ADP) in the apo includes residues in C4 and other suboptimal path include residues in C1 and C2 (Figure 8B; Supplemental Table 5). In the presence of substrates Asp13 communicates through N6-helix, Linker, and C4-helix residues. Alternative routes include residues in N3-helix, but are less statistically populated.

**Figure 8:**
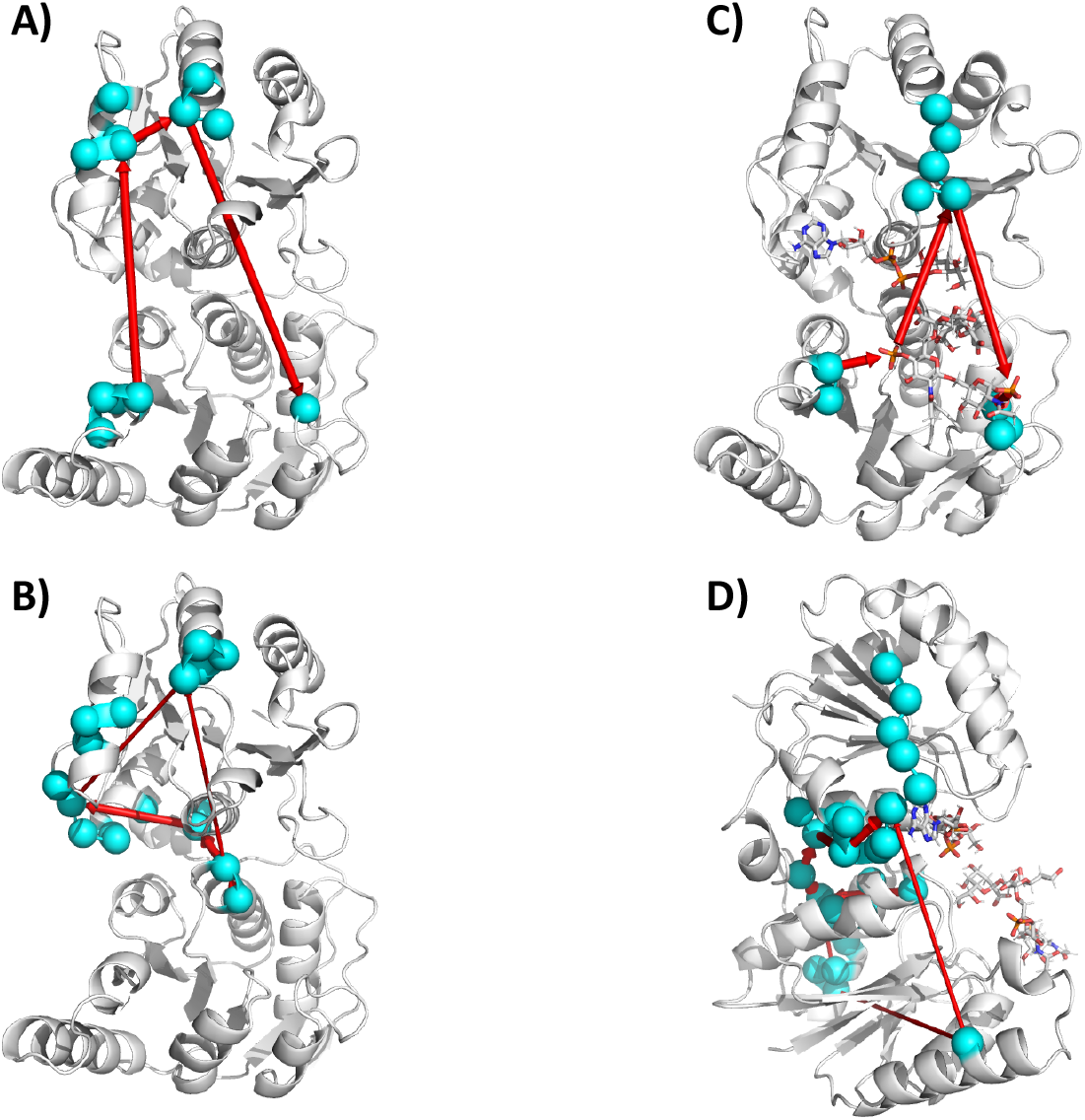
Protein communication network between residues Arg60/Arg120 and Asp13/Met242 for (**A**) HepI apo (Arg60/Arg120), (**B**) HepI apo (Asp13/Met242), (**C**) HepI with ADP-Hep and FDLA (Arg60/Arg120 and Asp13/Met242) and (**D**) HepI with ADP-Hep and FDLA (Arg60/Arg120 and Asp13/Met242).

## Discussion

Heptosyltransferase I from *E. coli* currently has four solved crystal structures with resolutions less than 2.4 Å. The structures consist of one apo (2GT1), one binary complex with the donor product ADP (2H1F), a binary complex with a fluorinated heptose donor analogue (2H1H), and a pseudo-ternary complex with a deacylated acceptor and non-hydrolyzable glycoside analogue of the donor (6DFE).^[18, 22]^ Multiple experimental studies have also been performed with HepI, which indicate that the protein binds substrates via a random bi-bi mechanism - where either substrate can bind to HepI, followed by binding of the other substrate.^[41]^ No experimental evidence exists to describe whether product release is ordered. Since HepI is involved in the LPS biosynthetic pathway and utilizes a membrane anchored substrate (Kdo_2_-Lipid A) in conjunction with a cytosolic nucleotide diphosphate sugar (ADP-Hep), HepI is expected to localize on the membrane to catalyze the transfer of the heptose sugar onto the Kdo_2_-Lipid A (Figure 1). Due to the soluble nature of ADP-Hep and its availability in the cytosol, HepI and ADP-Hep are anticipated to encounter one another prior to membrane localization. In addition, Kdo_2_-Lipid A induces a conformational change that would catalytically be unproductive in the absence of ADP-Hep. Therefore, ADP-Hep is anticipated to bind to HepI prior to the Kdo_2_-Lipid A. However in an effort to be fully rigorous, we simulated all possible binary and ternary complexes possible on the path for catalysis so that we could examine their significance and possible contribution (Figure 4). From the MMPBSA analysis, the binding free energy of ADP-Hep to HepI is approximately 3 kcal/mol lower than the binding free energy of Kdo_2_-Lipid A to HepI, which supports that the ADPHep•HepI binary complex is first to form because it is more energetically favorable. The binding of Kdo_2_-Lipid A to the binary complex of ADP-Hep•HepI is −8.75 kcal/mol. This is approximately 4 kcal/mol higher than the formation of the ADP-Hep ternary complex from the HepI and Kdo_2_-Lipid A binary complex. We hypothesize that the lower affinity of the Kdo_2_-Lipid A to the HepI and ADP-Hep binary complex is compensated for by the localization HepI and Kdo_2_-Lipid A to the membrane to facilitate this interaction.

### Effect of Substrate/Products on Local Dynamics

The backbone RMSD of all the simulations converge at approximately 2 Å and are relatively stable, even when extended for an additional 50 nanoseconds (Figure 2A). There are no obvious large-scale rearrangements that occur during the simulations, suggesting that the dynamics of HepI in the timescale of our simulations seems to be limited to local secondary structures. There are modest differences in the RMSF of the ternary substrate complex relative to the apo and are more clearly visible in the difference relative to apo (Figure 2B & 2C). Some of these differences occur in N-terminal residues, in the 60s (N3), 100s (N4), the linker (158-172), and 300s (C5). The N3 and N4 helices include arginine and lysine residues that have previously been shown to form electrostatic interactions with the acceptor phosphates as anchors in the N-terminal acceptor binding site. These interactions between the phosphates of FDLA and the basic residues of HepI in the N3/N4/N5 are present in our simulations (Figure 3A). The flexibility of N3 and N4 in the presence of the substrates relative to apo does not change, but in the presence of the products there is an increase in fluctuations and the standard deviation. The increase in dynamics of this region may be promoting product release. The N4 in both the presence of either the substrates or the products, increases in the local fluctuations and there are lysine residues that are transiently bound to the phosphates of the acceptor. In the linker region, both at the 150s and 200s there is an increase in dynamics which correspond to the loops that are connected to each of the respective domains. These regions may be responding to communication occurring between domains in the presence of substrate/products. The C4 and C5 are adjacent N3 and N4. In addition, C4 and C5 are adjacent to the hydrophobic pocket where the reaction may occur.

### Effect of Substrate/Products on Global Dynamics

Changes in the pK_a_ of Asp13 is consistent with previous mutagenesis and its implicated role as a catalytic base for HepI, with analogous Asp residues being conserved in all glycosyltransferases of the GT-B structural class (Supplemental Table 3).^[18]^ The hydrophobic environment and the increase in local negative charge from binding both substrates facilitates this rise in pK_a_. The shift for Asp13 is observed in the presence of the N-terminal substrate or product, r FDLA/FDHLA, is largely unchanged by binding of substrate or product to the C-terminal domain. The largest pK_a_ shift of Asp13 occurs in the presence of FDHLA in the binary and ternary complex. All of this points to the drastic effect of the sugar residues in close proximity to Asp13 driving up the pK_a_. This is consistent with observations in other systems, where changes in pK_a_ of buried acidic residues and the hydrophobic contribution of saccharide binding has been well documented and discussed.^[68, 69]^ In addition, the drop in pK_a_ of the adjacent lysine suggests further establishes the importance of this pocket becoming uncharged upon ligand complexation. We hypothesize that Lys7 may acts as a proton shuttle by abstracting a proton from the protonated Asp13, increasing the overall charge in the N-terminal domain, to facilitate product release, as seen by the instability of ADP, and to a lesser extent FDHLA, in the active site in the absence of restraints when Asp13 is deprotonated. Furthermore, the release of ADP by the FDHLA•HepI complex as, opposed to the release of FDHLA by the ADP•HepI, is supported by the more negative binding free energy of FDHLA to the ADP•HepI complex.

GT-Bs are expected to undergo a global conformational change prior to catalysis. The distance between the center of masses between each domain was used measured over time. In Figure 3, the apo enzyme has a center of mass distance centered around 30.5 Å, whereas the ternary complex with the substrates is centered closer to 31.0 Å and, the ternary complex with the products is centered around 29.5 Å. This suggests that in the presence of the substrates, HepI prefers a more “open” conformation and in the presence of the products it prefers a more “closed” conformation relative to the apo. Furthermore, the protonated Asp13 product complex has a 28.5 Å distance between the center of masses of the two domains. During the course of the simulation, FDHLA gets close enough to the C terminal domain that it begins to interact with residues Arg189 and Lys192, interactions that are not observed in any of the other complexes. Analysis of the hinge motion of the protein via Dyndom^[70, 71]^ shows an 86.6% closure when compared to the apo crystal structure (PDB:2GT1). We do not observe a full closing of the protein, however this is hypothesized to be a limitation of the timescale (100 nanosecond) utilized in this study.

From examination of the interdomain residue interactions, in the apo state the motion of the residues in the N3 helix are negatively correlated with residues in the C2 helix. These two helices are directly across the interdomain gap from each other and would require their motions to coordinate in a negatively correlated fashion to facilitate an “open” to “close” transition. These two helices may also be dynamic and correlated to promote substrate capture by each domain. As demonstrated in Figure 6, in the presence of the substrates/products the coupled motions between N3 and C2 is preserved relative to apo. This positively correlated motion may be indicative of substrate capture. This asymmetric pincer mode is dominant in PC1(apo) and PC2(substrates), but both only account for less than 20% of their respective contribution to the total variance (Figure 5, Supplemental Table 4). This transition from most dominant motion in the apo, to second dominant in the presence of the substrates suggests a transition from this substrate capture to a more catalytically productive mode. In the presence of the substrates there is an enhancement of positively correlated motions between N5/N6 and C5/C6 relative to the apo (Figure 6). This motion is relevant for the potential conformational rearrangement that occurs prior to catalysis and is evident in PC2/3 (apo) and PC1(substrates). In PC2/3(apo), the dynamics are still centered around N3 and C2, but an increase in the dynamics at N5/N6 and C5/C6 may be those low populated modes that contribute to catalysis post substrate binding, but these two combined only account for 25% of the variance. In contrast, PC1(substrates) the N5/N6 and C5/C6 have the greatest fluctuations and this could be a coordinated effort between domains to promote catalysis by moving antiparallel to one another in a twisting motion that is conducive to aligning the hydroxyl of the FDLA (nucleophile) and the anomeric carbon of the ADP-Hep (electrophile) closer to the hydrophobic pocket. This mode alone accounts for 30.2% of the variance which speaks to the increased population of this state relative to the apo (Supplemental Table 4). The product complex has a greater scattered population of negatively correlated motions relative the substrate complex and apo (Figure 4). Upon protonation of Asp13 in the product ternary complex, this scattering is diminished and there is an enhancement of negatively correlated interdomain motions between N4 with C1/C4/C5/C6 and N5 with C1/C4/C5/C6 (Figure 6D). More interestingly, PC1 (ADP-Hep•FDLA•HepI (D13+H)) has the most fluctuations in the C terminal C3-C5 (Supplemental Figure 5). This motion brings the C terminal domain closer to the N-terminal domain in a more typical “closing” motion. This principal component accounts for 40.7% of the variance alone (Supplemental Table 4). This, along with the changes in the center of mass distance strongly suggest this may be the beginning of the global conformational change this enzyme undergoes prior to catalysis.

The communications pathways we observe within and between domains both in the presence and absence of substrates provides a mechanism in which catalysis can be facilitated by substrate binding. In the apo state, communication between residues Arg60 and Arg120, which were previously determined to be important for FDLA binding^[40]^ form shortest paths through C1- and C2-helix residues (Figure 8A; Supplemental Table 5; Supplemental Figure 9A). This intradomain communication is most likely mediated by electrostatic interactions between positively charged residues of Arg60/Arg120 in N3/N5 and the negatively charged residues Glu224 in C2, and Glu196 and Glu197 in C1. Interestingly, Trp62 is also involved in this communication network, and this has previously been hypothesized to act as a local reporter for FDLA binding.^[39]^ Communication with this Trp62 residue is lost in the presence of substrates and products in the “open” (deprotonated Asp13) and partially “closed” (protonated Asp13) states which suggests this residue undergoes a rearrangement which “uncouples” it from this communication network (Figure 8C; Supplemental Table 5; Supplemental Figure 9D). This is consistent with Trp62 acting as a local reporter for FDLA binding. In the partially “closed” product state (protonated Asp13), communication between Arg60/Arg120 involves Trp199 which was shown to be a major contributor to HepI tryptophan fluorescence blue shift in the presence of the acceptor (Supplemental Table 5).^[39]^ Trp199 is in the C terminal domain and this network is only seen in the partially “closed” state which shows that Trp199, unlike Trp62, acts not as a reporter for substrate binding, rather as a reporter for the coordinated “closing” motion that occurs prior to catalysis. Pathways between Asp13 (the catalytic base) and the Met242, which is important for hydrogen bonding with the adenine ring of ADPHep, also involves Trp199 in the apo (Figure 8B; Supplemental Table 5). In the presence of substrates, communication with 199 is lost and communicates through residues in the linker (Figure 8D; Supplemental Table 5). The disruption of communication between N1 and C1 may setup Trp199 to now communicate with Arg60/Arg120 to go from substrate search to undergo the conformational change required for catalysis. Furthermore, communication between Asp13/Met242 through the linker is present in the presence of the substrates in the “open” state, but disappears in the presence of the products in the partially “closed” state (Figure 8D; Supplemental Table 5). Dyndom analysis suggests residues 163-165 are the fulcrum in which the two domains bend towards one another. Tyr163 is one of these hinge residues and this communication network and could be the post substrate binding precursor to the global conformational change. Once the substrates bind triggering rearrangement of N and C terminal residues, these contacts facilitate a network of rearrangements in both domains causing their closer to another. Communication through the linker is most likely lost as it moves away from the back side of two Rossman-domains creating a small back side pocket, which was previously implicated to be an allosteric binding site for small HepI inhibitory compounds,^[72]^ and because as HepI enters the partially “closed” product state, residues at the interface of the two domains are now able to make direct contacts with those across the interdomain gap.

### Conclusion

In conclusion, we have performed molecular dynamic simulations of HepI in substrate/product binary and ternary complexes to gain a better understanding of the dynamics that govern this family of proteins. BFE analysis allowed determination of the substrate binding order and product release order. In addition, we have begun to unravel the complex network of communication between domains that facilitates substrate binding, product release, and global conformational changes that lead to catalysis. These results support the hypothesis that the residues involved in ligand binding from each domain communicate ligand occupancy to the other ligand pocket, ensuring that the enzyme doesn’t undergo large closure events that would be unproductive in the absence of bound ligands. This work provides insight that may be useful towards the design of new inhibitors against the Heptosyltransferase family of proteins, and also other GT-B enzymes.

## Abbreviations

GT: glycosyltransferase
CAZY: Carbohydrate-Active enZYmes Database
HepI: Heptosyltransferase I
ADP-Hep: ADP-L-glycero-β-D-manno-heptose
Kdo: 3-deoxy-D-*manno*-oct-2-ulosonic acid
H-Kdo_2_-Lipid A: heptosylated Kdo_2_-Lipid A
FDLA: fully-deacylated Kdo_2_-Lipid A
ADP: adenosine disphosphate
FDHLA: fully-deacylated heptosylated-Kdo_2_-Lipid A
FDLA-H: deprotonated sugar donor nucleophile
D13+H: protonated aspartic acid 13
TIP3P: transferrable intermolecular potential with 3 points
RMSD: root mean square deviation
R_gyr_: radius of gyration
RMSF: root mean square fluctuations
PCA: Principal component analysis
DCCM: dynamic cross correlation matrix
MMPBSA: molecular mechanics Poisson Boltzmann solvent accessibility

## Acknowledgement

Kelly Thayer for providing tutorials to allow sector analysis. This work was supported by a grant from the National Institutes of Health (1R15AI119907-01).

## TOC Graphic

## Supplemental

**Supplemental Table 1:**
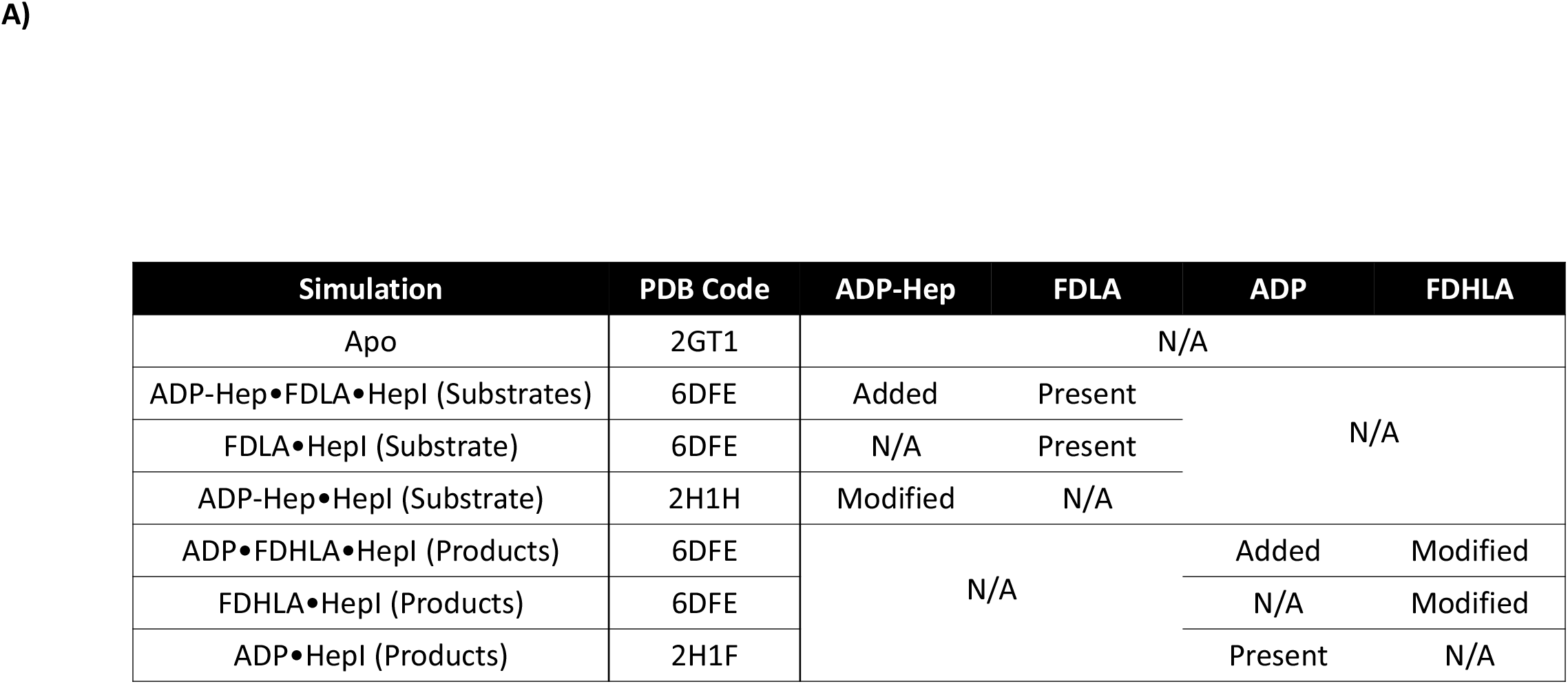
The crystal structures used for simulations and details on whether a ligand was present in the original structure or if it was added. The fluorinated ADP-Hep was taken from PDB:2H1H and the fluorine was replaced with a hydroxyl group in the proper stereoconfiguration. The FDHLA was constructed by the addition of a Heptose on the FDLA in the right stereo/regioconfiguration.

**Supplemental Table 2:**
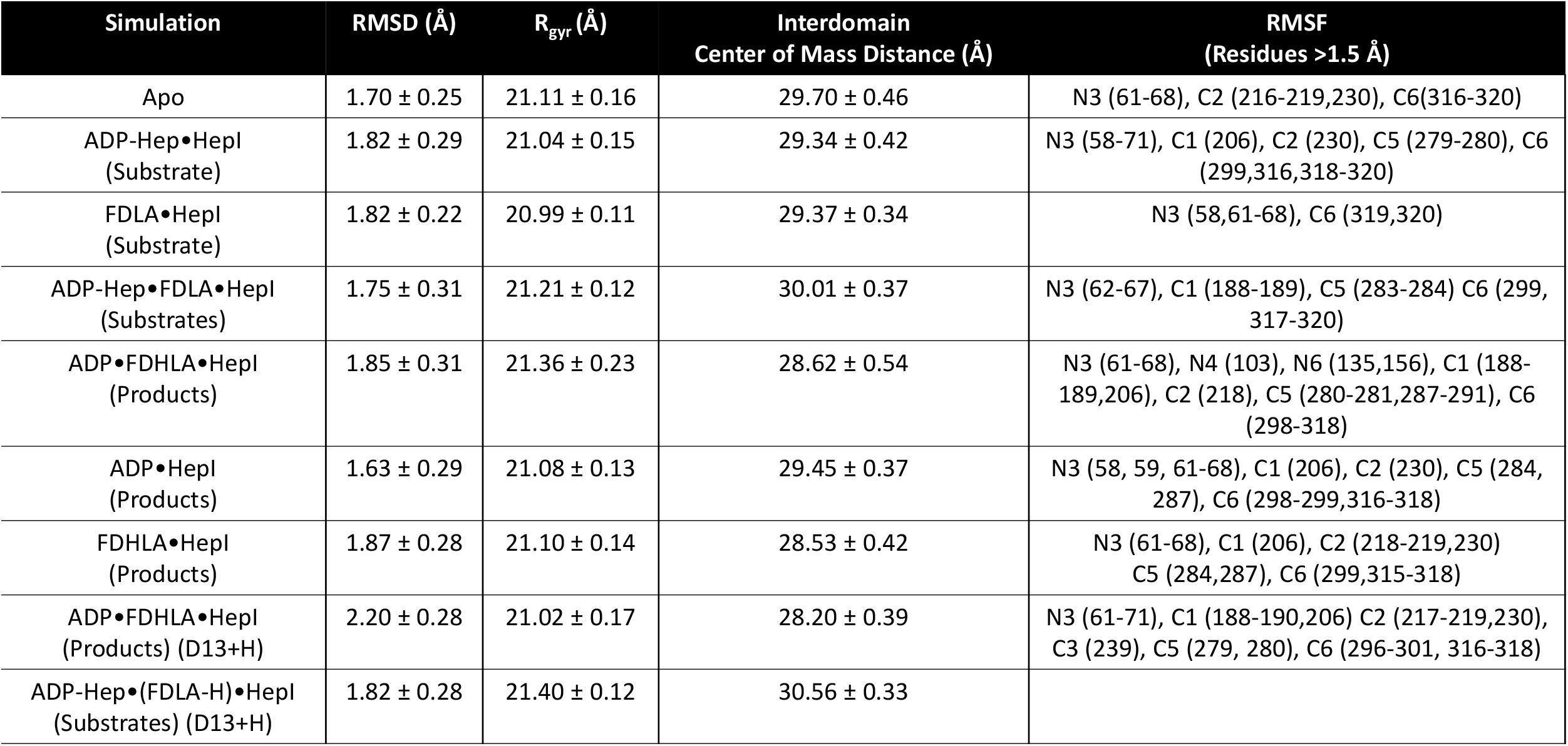
Average values of RMSD and radius of gyration from the length of one representative simulation with standard deviations. RMSF values are reported for regions and residues that have a greater than 1.5 Å fluctuation.

**Supplemental Table 3:**
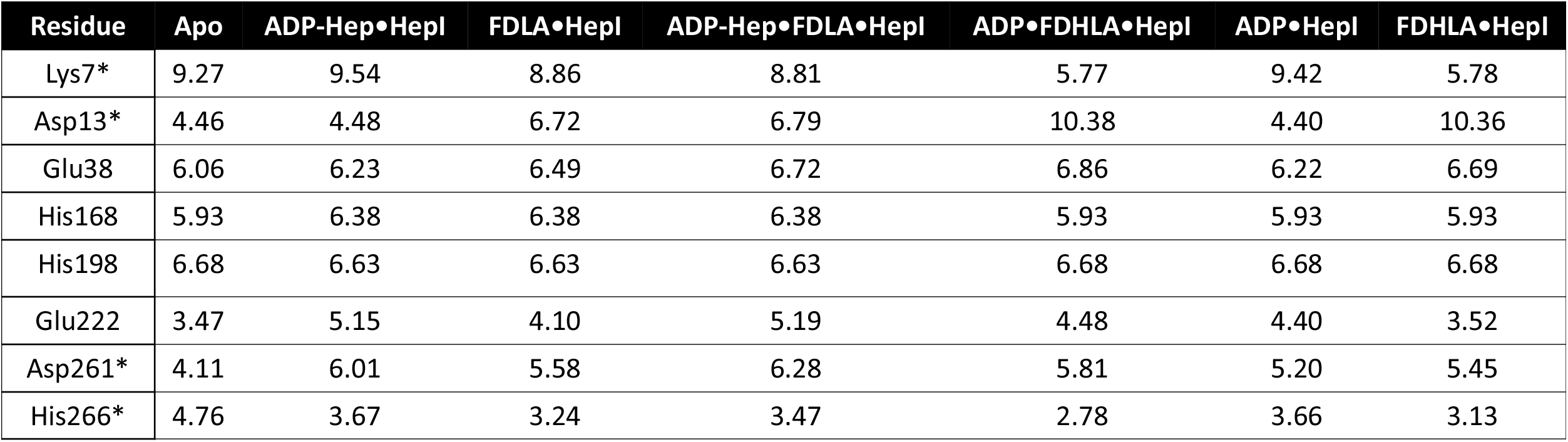
pK_a_ of ionizable sidechains as determined by PROPKA3.

**Supplemental Table 4:**
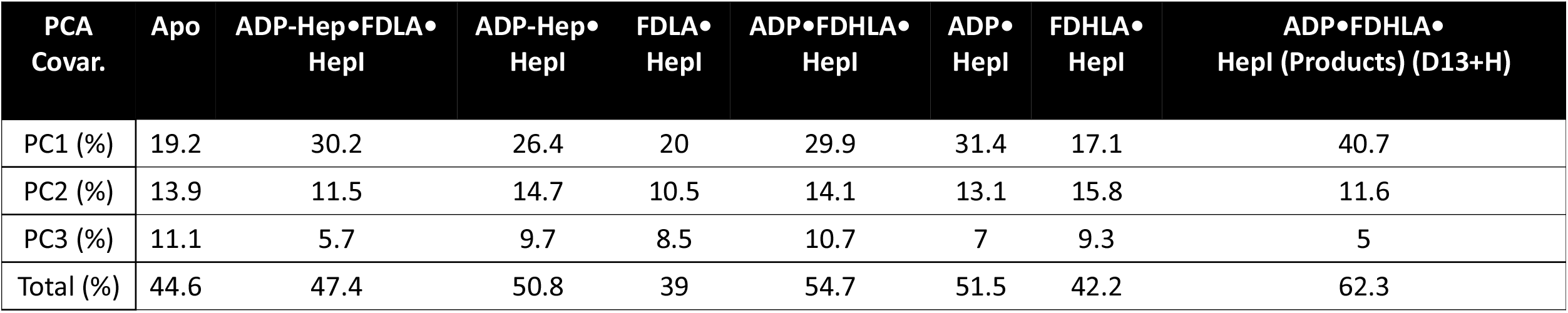
PCA percent covariance table.

**Supplemental Table 5:**
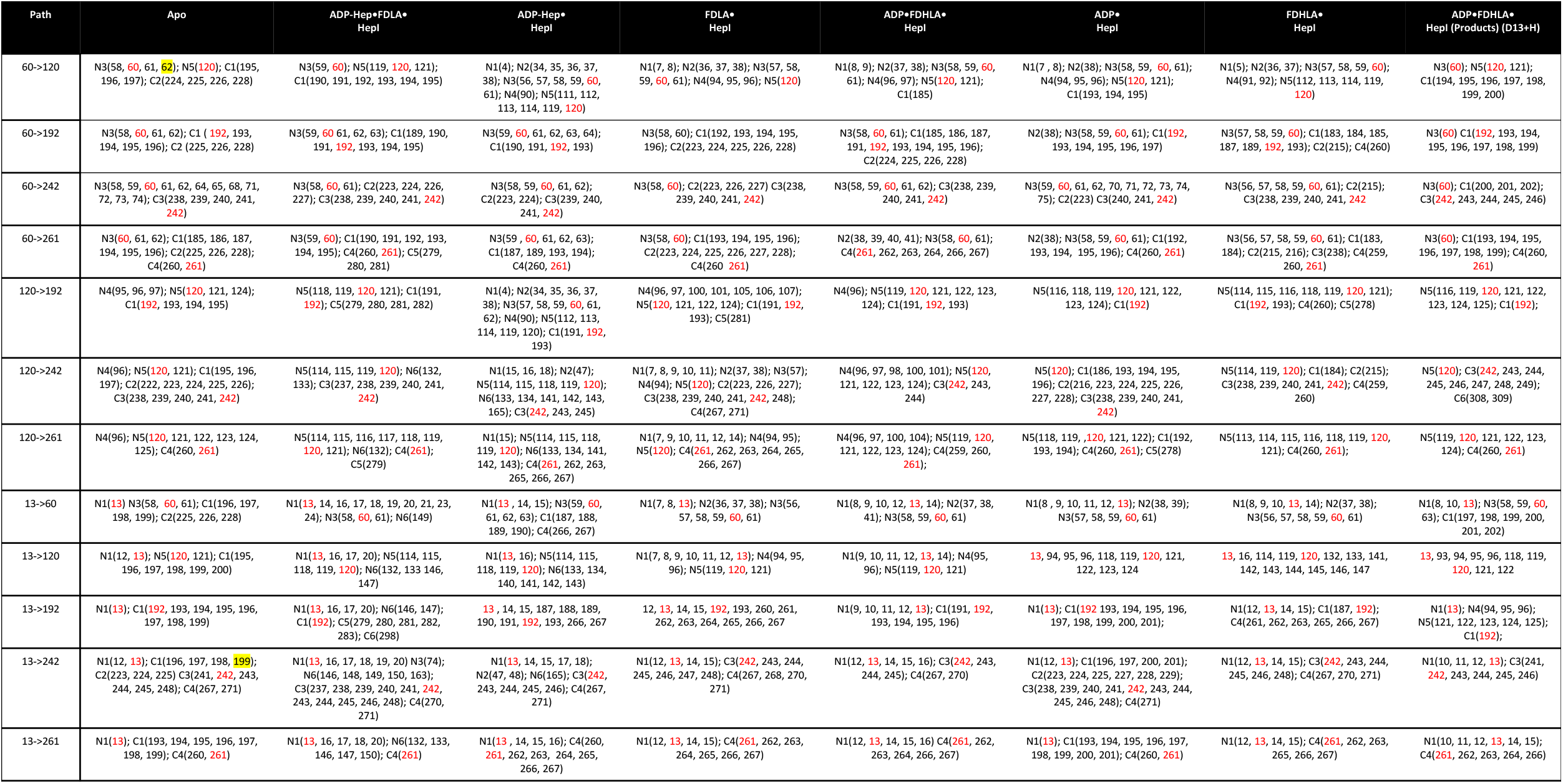
Consensus residues involved in each of the collection of shortest paths of communication in their respective simulations.

**Supplemental Figure 1:**
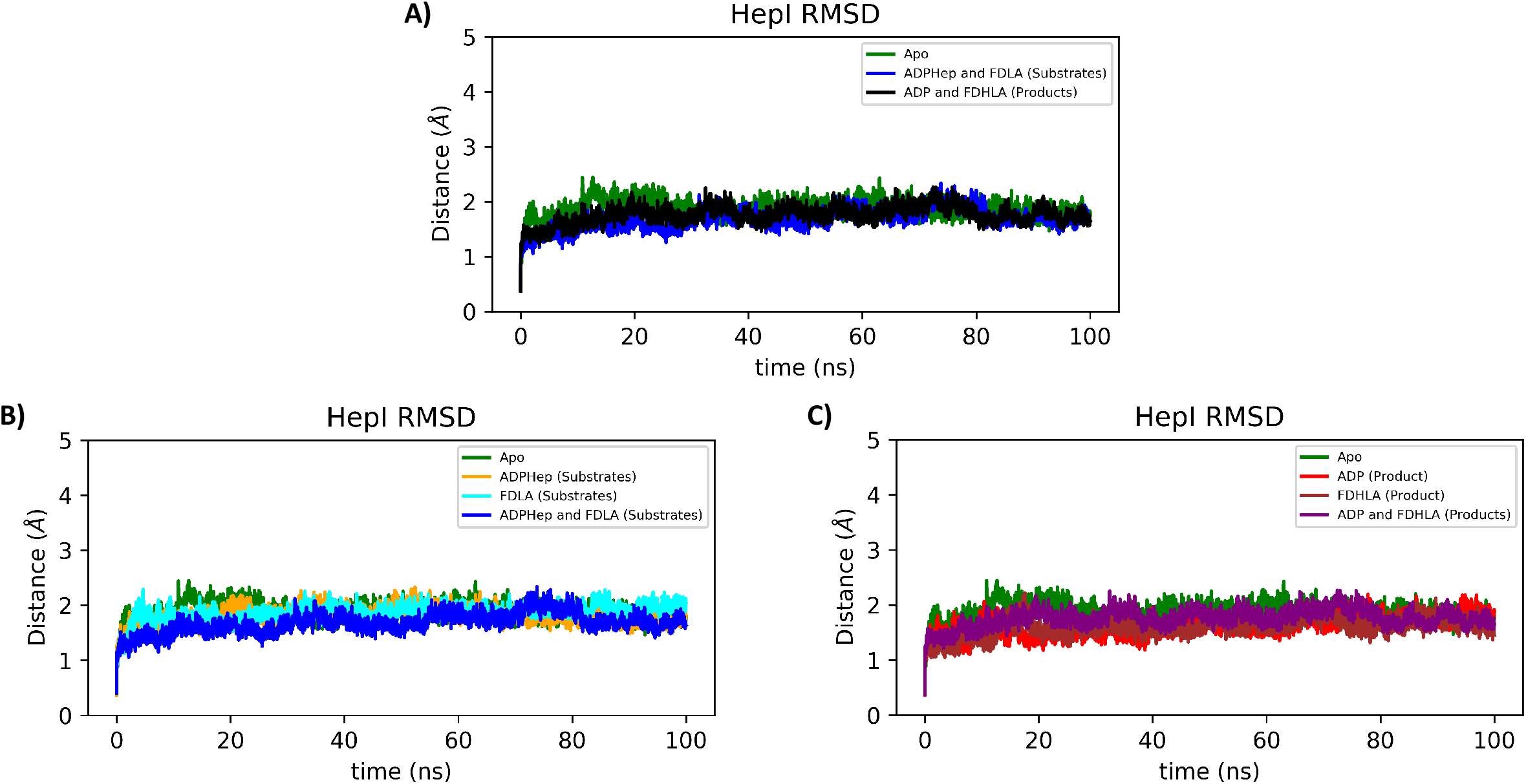
(**A**) Backbone RMSD of HepI apo, substrate ternary complex and product ternary complex with deprotonated Asp13. (**B**) Backbone RMSD of HepI apo, substrate binary complexes (ADP-Hep•HepI, FDLA•HepI) and substrate ternary complex. (**C**) Backbone RMSD of HepI apo, product binary complexes (ADP•HepI, FDHLA•HepI) and product ternary complex with deprotonated Asp13.

**Supplemental Figure 2:**
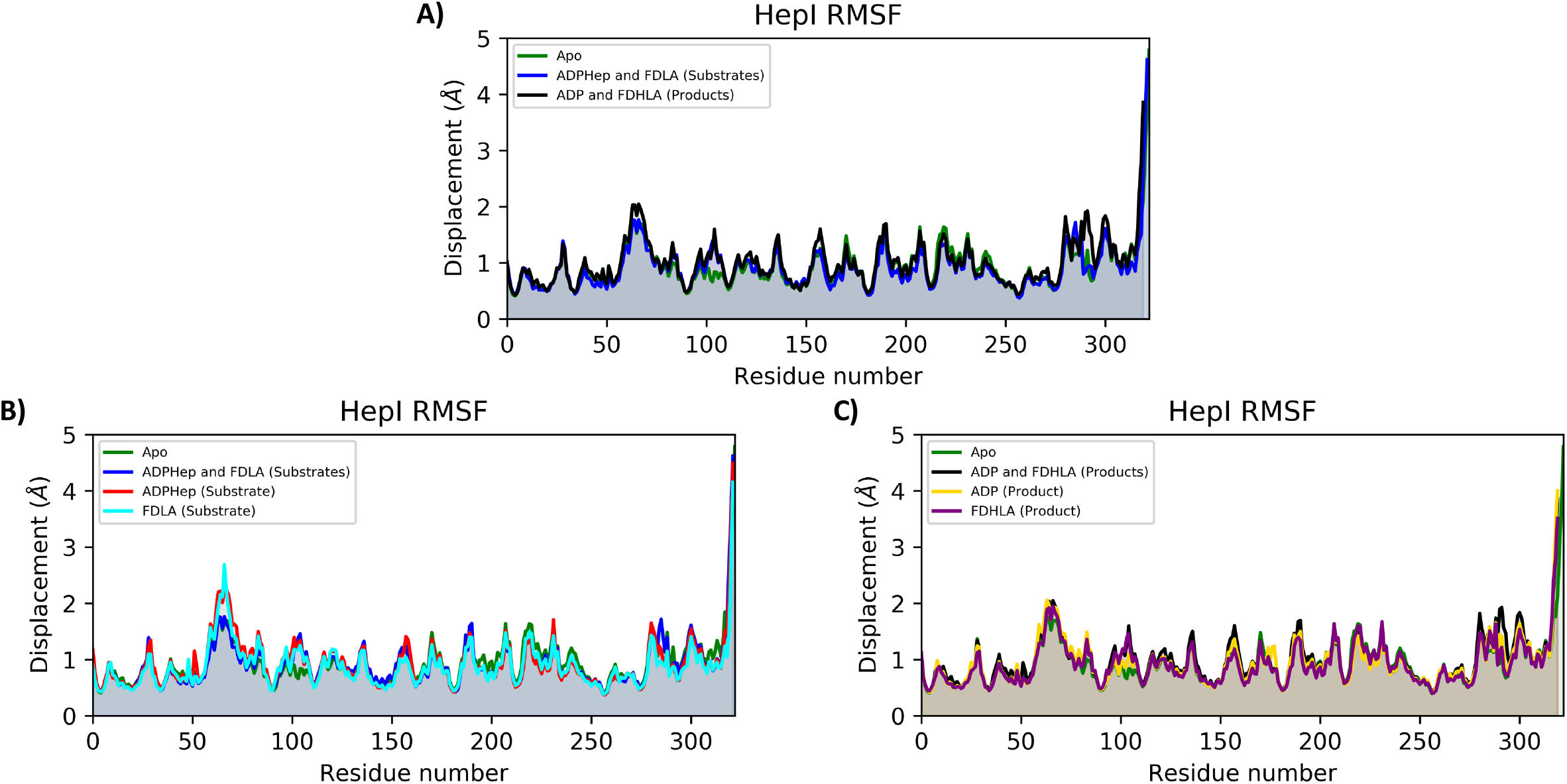
(**A**) C_α_ RMSF of HepI apo, substrate ternary complex and product ternary complex with deprotonated Asp13. (**B**) C_α_ RMSF of HepI apo, substrate ternary complex and substrate binary complexes (ADP-Hep•HepI, FDLA•HepI). (**C**) C_α_ RMSF of HepI apo, product ternary complex with deprotonated Asp13 and product binary complexes (ADP•HepI, FDHLA•HepI).

**Supplemental Figure 3:**
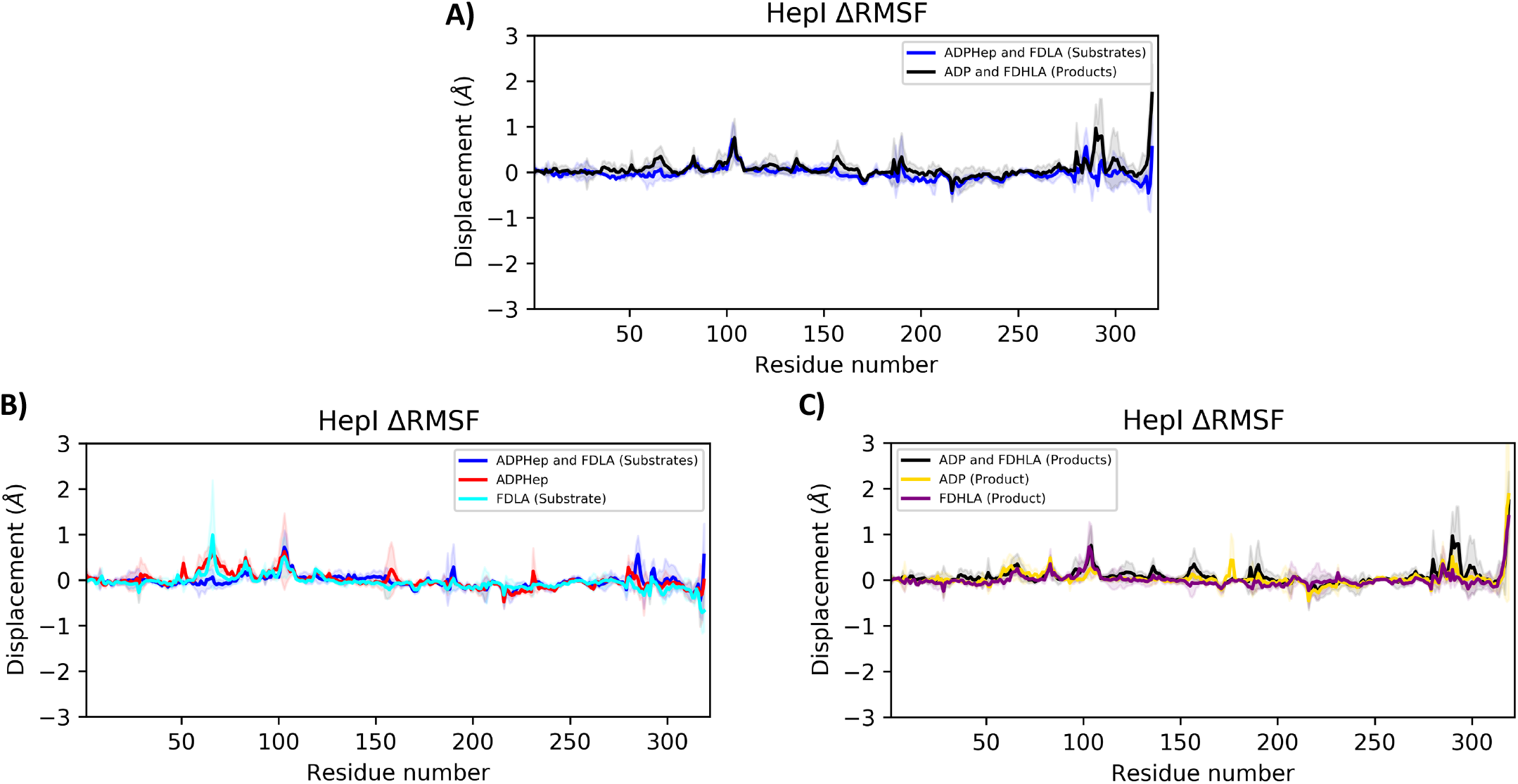
(**A**) C_α_ ΔRMSF of HepI substrate ternary complex and product ternary complex with deprotonated Asp13. (**B**) C_α_ ΔRMSF of HepI substrate ternary complex and substrate binary complexes (ADP-Hep•HepI, FDLA•HepI). (**C**) C_α_ΔRMSF of HepI product ternary complex with deprotonated Asp13 and product binary complexes (ADP•HepI, FDHLA•HepI). solid lines are average differences relative to apo (i.e. RMSF_substrates_-RMSF_apo_) and shaded region is the standard deviation of the average difference. Positive values indicate those residues are more flexible relative to HepI apo.

**Supplemental Figure 4:**
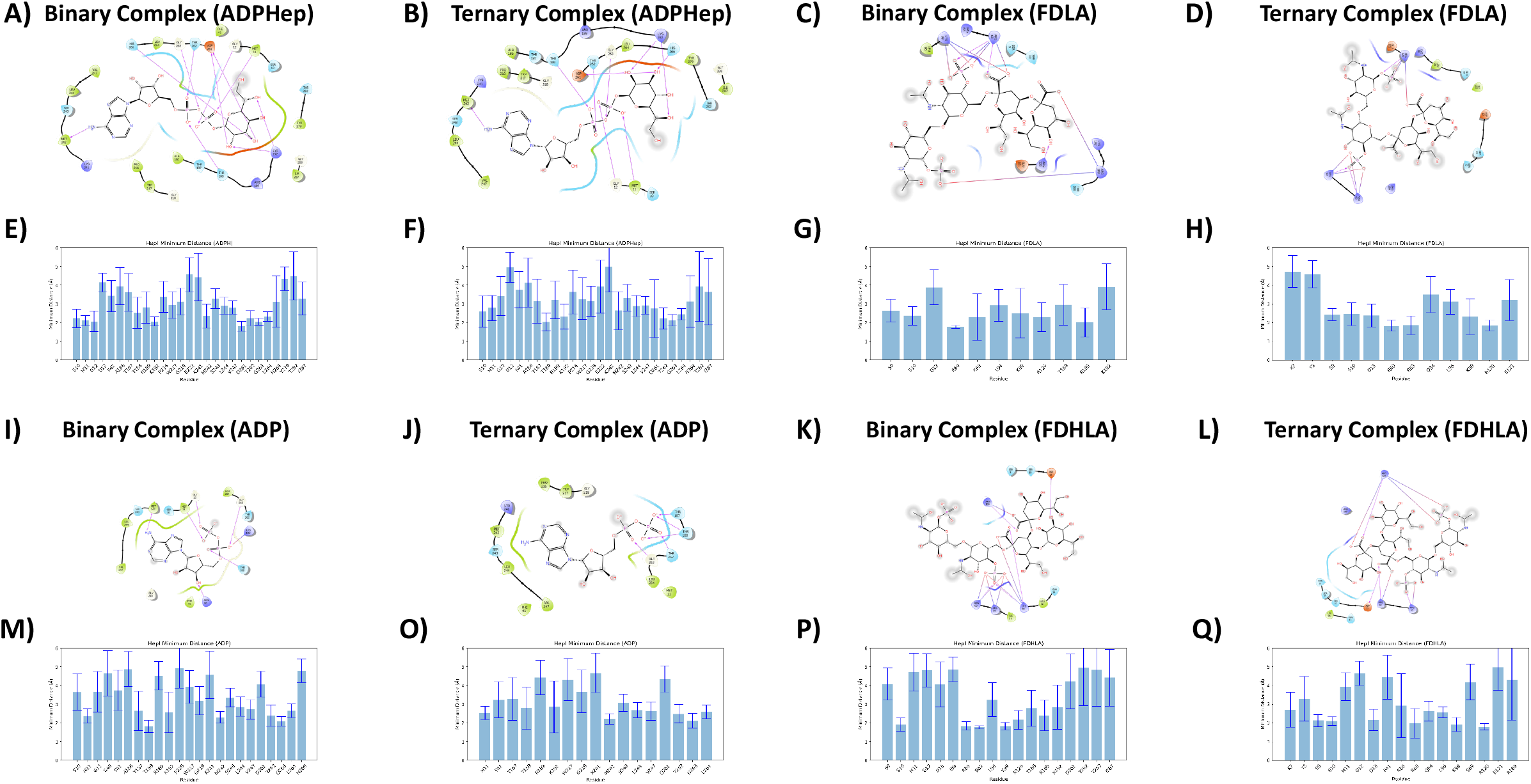
Ligand interaction diagram of (**A-B**) ADP-Hep, (**B-C**) FDLA, (**I-J**) ADP and (**K-L**) FDHLA from binary and ternary complexes. Bar plots of residues with average minimum distances of less than 5 Å from (**E-F**) ADP-Hep, (**G-H**) FDLA, (**M-O**) ADP and (**P-Q**) FDHLA from binary and ternary complexes.

**Supplemental Figure 5:**
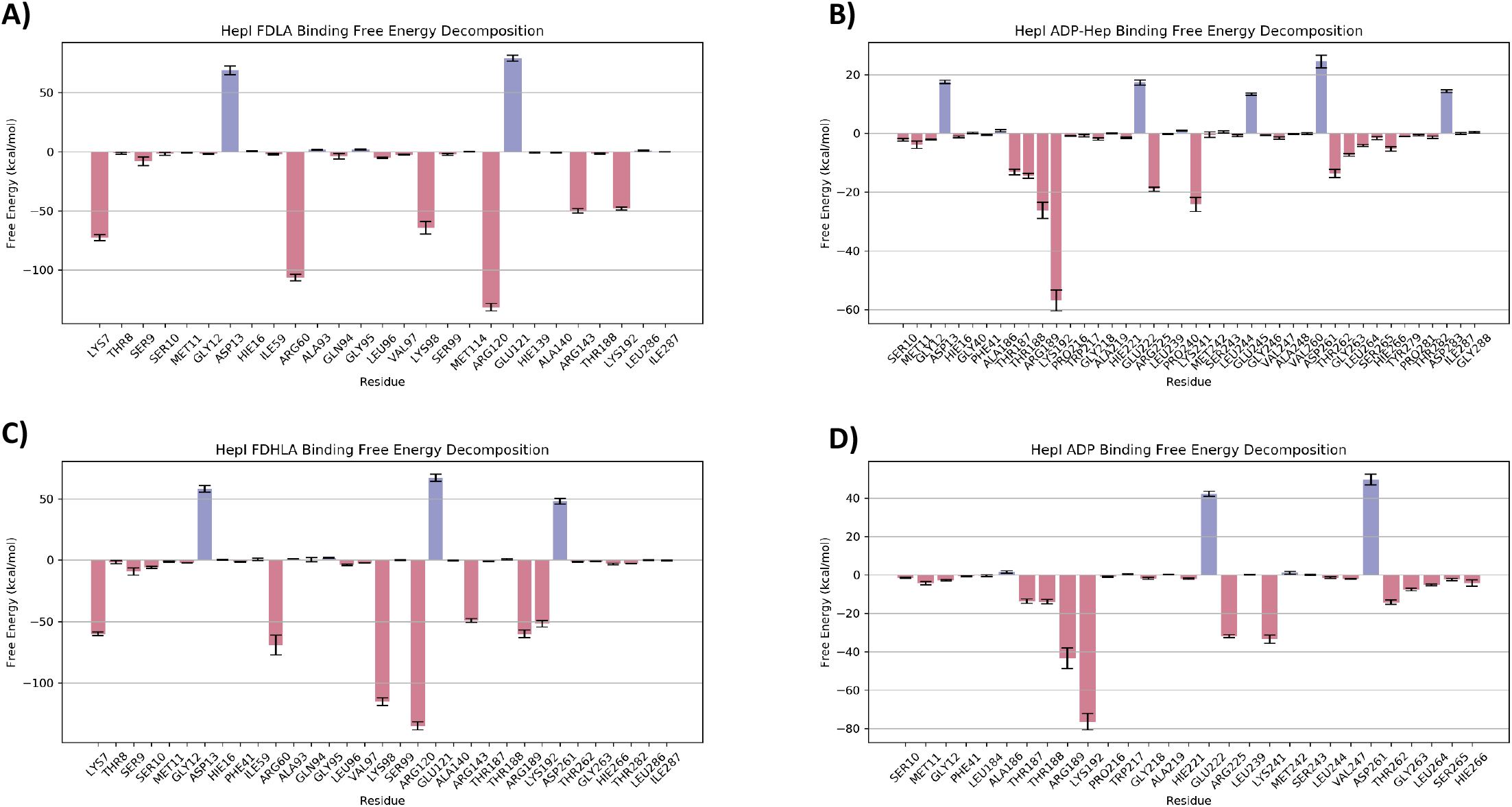
Binding free energy contribution of residues within 6 Å of (**A**) FDLA, (**B**) ADP-Hep, (**C**) FDHLA and (**D**) ADP in the ternary complex.

**Supplemental Figure 6:**
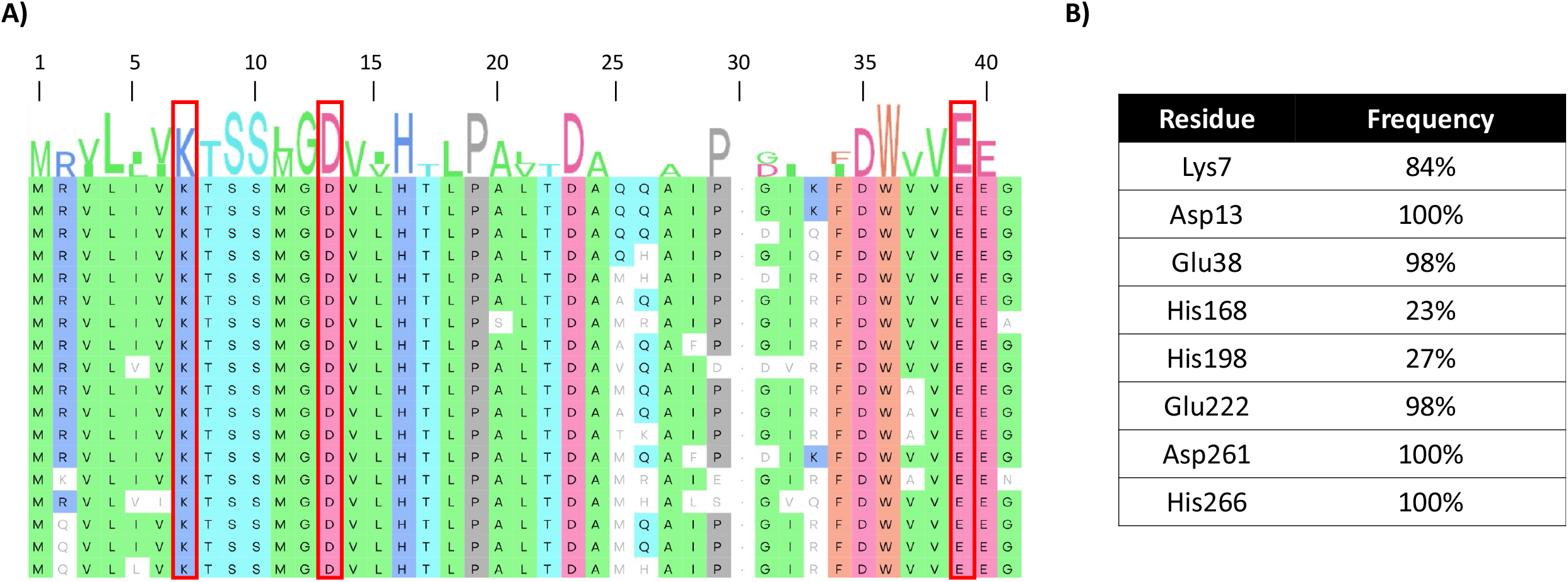
(**A**) 18 representative sequences of Heptosyltransferase I from a multiple sequence alignment of 150 sequences from various homologues with residues highlighted in red boxes that demonstrate a large pK_a_ shift. (**B**) Table of residues identified with pK_a_ shifts and their corresponding frequency of occurrence at their respective positions in the multiple sequence alignment.

**Supplemental Figure 7:**
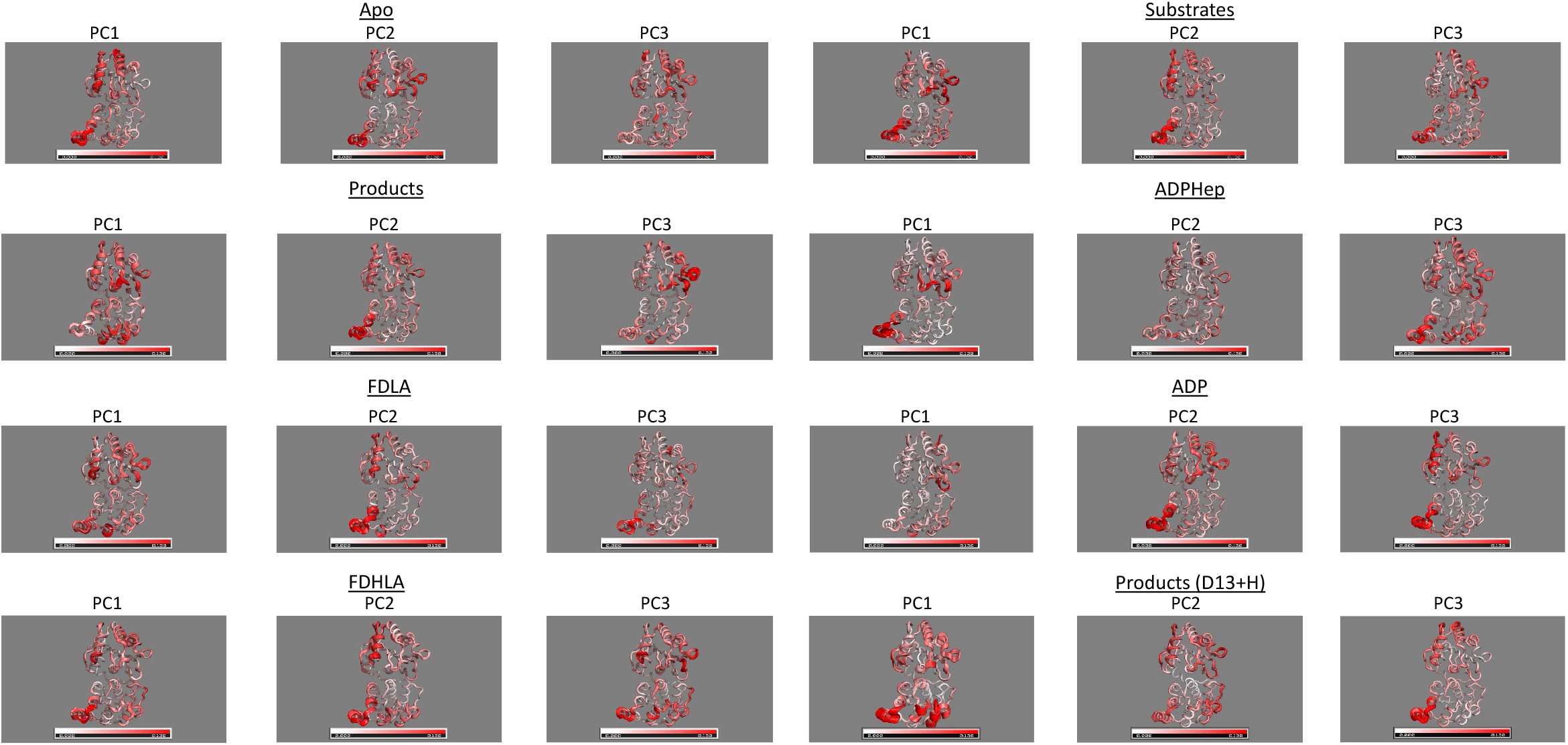
First three principle components of (**A**) HepI apo and (**B**)) substrate complex (**C**)) product complex (**D**) ADP-Hep binary complex (**E**) FDLA binary complex (**F**) ADP binary complex (**G**) FDHLA binary complex and (**H**) product ternary complex with protonated Asp13.

**Supplemental Figure 8:**
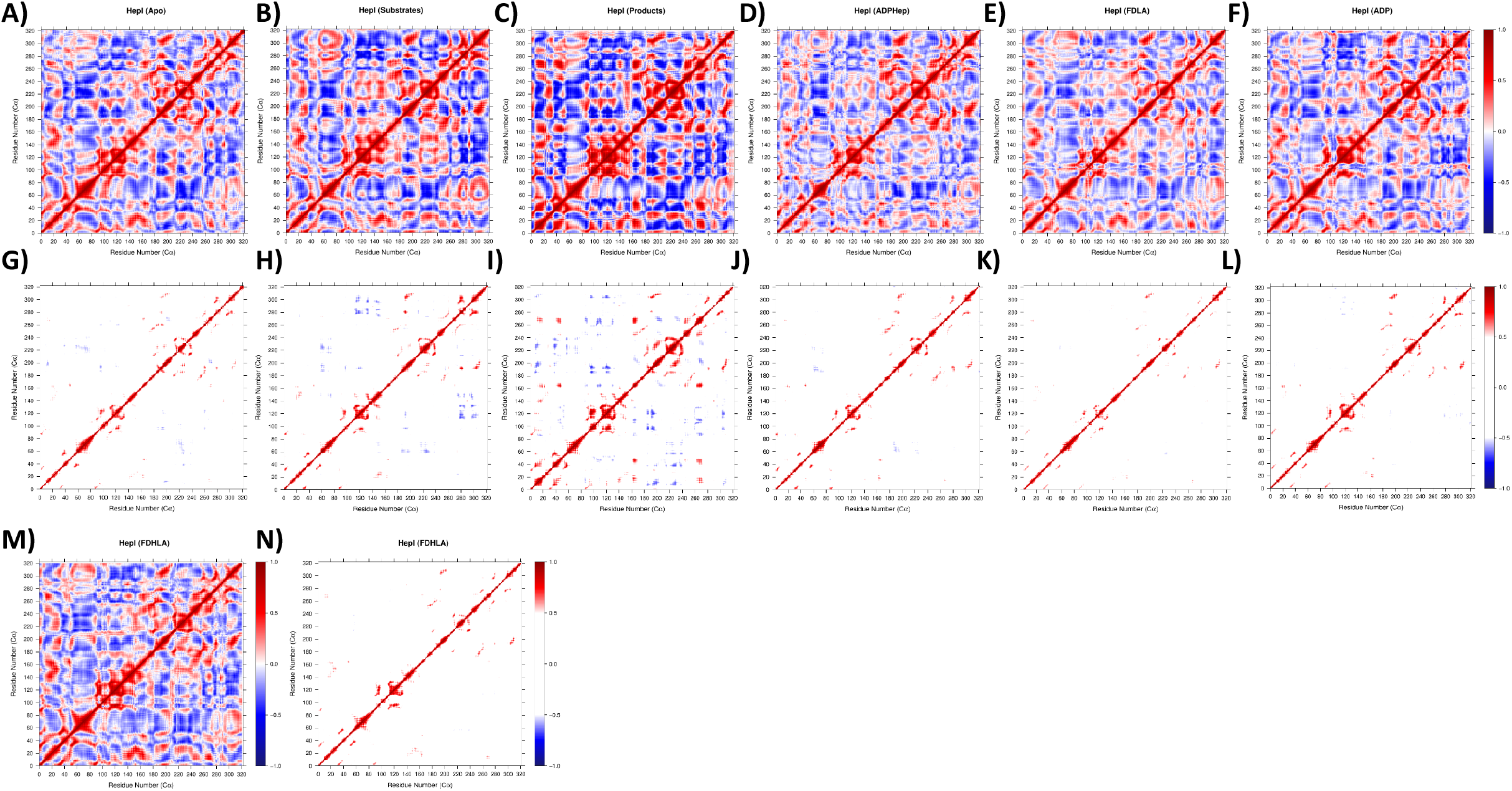
Dynamic cross correlation map of (**A,G**) HepI apo, (**B,H**) ADP-Hep•FDLA•HepI, (**C,I**) ADP•FDHLA•HepI, (**D,J**) ADP-Hep•HepI, (**E,K**) FDLA•HepI, (**F,L**) ADP•HepI, and (**M,N**) FDHLA•HepI.

**Supplemental Figure 9:**
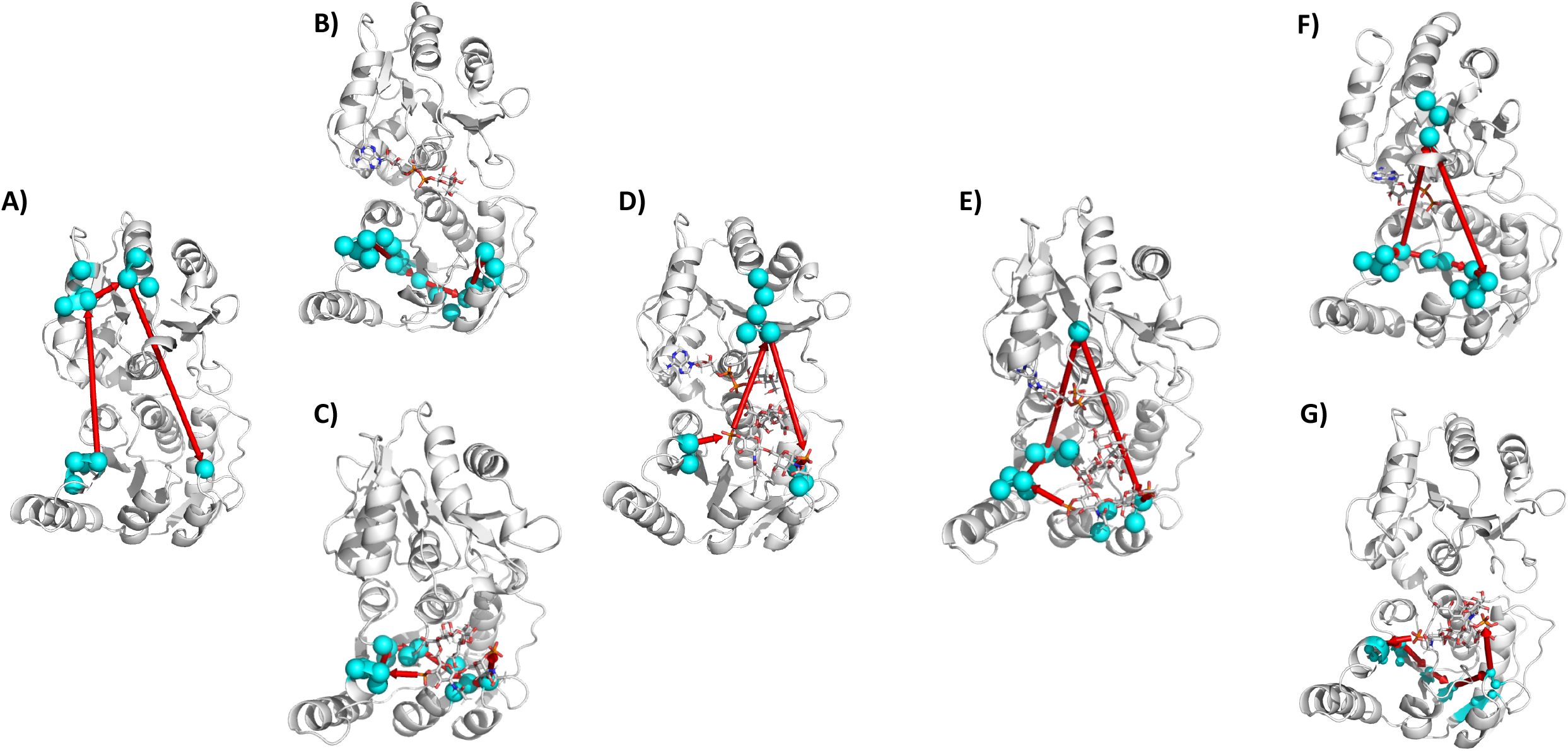
Protein communication network between residues Arg6O and Arg120 for (**A**) HepI apo, (**B**) with ADP-Hep, (**C**) with FDLA, (**D**) with both substrates, (**E**) with both products and deprotonated Asp13, (**F**) with ADP and (**G**) with FDHLA.

## References

1. Hamilos, D.L., Biofilm Formations in Pediatric Respiratory Tract Infection Part 2: Mucosal Biofilm Formation by Respiratory Pathogens and Current and Future Therapeutic Strategies to Inhibit Biofilm Formation or Eradicate Established Biofilm. Curr Infect Dis Rep, 2019. 21(2): p. 8.

2. Rosenbalm, K.E., et al., Glycomics-informed glycoproteomic analysis of site-specific glycosylation for SARS-CoV-2 spike protein. STAR Protoc, 2020. 1(3): p. 100214.

3. Lenza, M.P., et al., Structural Characterization of N-Linked Glycans in the Receptor Binding Domain of the SARS-CoV-2 Spike Protein and their Interactions with Human Lectins. Angew Chem Int Ed Engl, 2020. 59(52): p. 23763–23771.

4. Elbatrawy, A.A., E.J. Kim, and G. Nam, O-GlcNAcase: Emerging Mechanism, Substrate Recognition and Small-Molecule Inhibitors. ChemMedChem, 2020. 15(14): p. 1244–1257.

5. Vajaria, B.N. and P.S. Patel, Glycosylation: a hallmark of cancer? Glycoconj J, 2017. 34(2): p. 147–156.

6. Schuman, B., J.A. Alfaro, and S.V. Evans, Glycosyltransferase Structure and Function, in Bioactive Conformation I, T. Peters, Editor. 2007, Springer Berlin Heidelberg: Berlin, Heidelberg. p. 217–257.

7. Roychoudhury, R. and N.L. Pohl, New structures, chemical functions, and inhibitors for glycosyltransferases. Curr Opin Chem Biol, 2010. 14(2): p. 168–73.

8. Lombard, V., et al., The carbohydrate-active enzymes database (CAZy) in 2013. Nucleic Acids Res, 2014. 42(Database issue): p. D490–5.

9. Chang, A., et al., Glycosyltransferase structural biology and its role in the design of catalysts for glycosylation. Curr Opin Biotechnol, 2011. 22(6): p. 800–8.

10. Breton, C., et al., Structures and mechanisms of glycosyltransferases. Glycobiology, 2006. 16(2): p. 29R–37R.

11. Breton, C., S. Fournel-Gigleux, and M.M. Palcic, Recent structures, evolution and mechanisms of glycosyltransferases. Curr Opin Struct Biol, 2012. 22(5): p. 540–9.

12. Montgomery, A.P., et al., Computational Glycobiology: Mechanistic Studies of Carbohydrate-Active Enzymes and Implication for Inhibitor Design. Adv Protein Chem Struct Biol, 2017. 109: p. 25–76.

13. Fan, F., et al., Structures and mechanisms of the mycothiol biosynthetic enzymes. Curr Opin Chem Biol, 2009. 13(4): p. 451–9.

14. Ardevol, A., et al., The reaction mechanism of retaining glycosyltransferases. Biochem Soc Trans, 2016. 44(1): p. 51–60.

15. Lee, S.S., et al., Mechanistic evidence for a front-side, SNi-type reaction in a retaining glycosyltransferase. Nat Chem Biol, 2011. 7(9): p. 631–8.

16. Winchell, K.R., et al., A Structural, Functional, and Computational Analysis of BshA, the First Enzyme in the Bacillithiol Biosynthesis Pathway. Biochemistry, 2016. 55(33): p. 4654–65.

17. Gloster, T.M., Advances in understanding glycosyltransferases from a structural perspective. Curr Opin Struct Biol, 2014. 28: p. 131–41.

18. Grizot, S., et al., Structure of the Escherichia coli heptosyltransferase WaaC: binary complexes with ADP and ADP-2-deoxy-2-fluoro heptose. J Mol Biol, 2006. 363(2): p. 383–94.

19. Li, Z., et al., Structural basis of Notch O-glucosylation and O-xylosylation by mammalian protein-O-glucosyltransferase 1 (POGLUT1). Nat Commun, 2017. 8(1): p. 185.

20. Volkers, G., et al., Structure of human ST8SiaIII sialyltransferase provides insight into cell-surface polysialylation. Nat Struct Mol Biol, 2015. 22(8): p. 627–35.

21. Brazier-Hicks, M., et al., Characterization and engineering of the bifunctional <em>N<em>-and <em>O<em>-glucosyltransferase involved in xenobiotic metabolism in plants. Proceedings of the National Academy of Sciences, 2007. 104: p. 20238–20243.

22. Blaukopf, M., et al., Insights into Heptosyltransferase I Catalysis and Inhibition through the Structure of Its Ternary Complex. Structure, 2018. 26(10): p. 1399–1407 e5.

23. Ramirez, A.S., et al., Structural basis of the molecular ruler mechanism of a bacterial glycosyltransferase. Nat Commun, 2018. 9(1): p. 445.

24. Ge, C., et al., Membrane remodeling capacity of a vesicle-inducing glycosyltransferase. FEBS J, 2014. 281(16): p. 3667–84.

25. Salinas, S.R., et al., Binding of the substrate UDP-glucuronic acid induces conformational changes in the xanthan gum glucuronosyltransferase. Protein Eng Des Sel, 2016. 29(6): p. 197–207.

26. Albesa-Jove, D., et al., Structural Snapshots and Loop Dynamics along the Catalytic Cycle of Glycosyltransferase GpgS. Structure, 2017. 25(7): p. 1034–1044 e3.

27. Snajdrova, L., et al., Molecular dynamics simulations of glycosyltransferase LgtC. Carbohydr Res, 2004. 339(5): p. 995–1006.

28. Righino, B., et al., Identification and Modeling of a GT-A Fold in the alpha-Dystroglycan Glycosylating Enzyme LARGE1. J Chem Inf Model, 2020. 60(6): p. 3145–3156.

29. Janos, P., et al., Three-dimensional homology model of GlcNAc-TV glycosyltransferase. Glycobiology, 2016. 26(7): p. 757–771.

30. Romero-Garcia, J., et al., Structure-function features of a Mycoplasma glycolipid synthase derived from structural data integration, molecular simulations, and mutational analysis. PLoS One, 2013. 8(12): p. e81990.

31. Panda, S.K., S. Saxena, and L. Guruprasad, Homology modeling, docking and structure-based virtual screening for new inhibitor identification of Klebsiella pneumoniae heptosyltransferase-III. J Biomol Struct Dyn, 2020. 38(7): p. 1887–1902.

32. Kadrmas, J.L. and C.R. Raetz, Enzymatic synthesis of lipopolysaccharide in Escherichia coli. Purification and properties of heptosyltransferase i. J Biol Chem, 1998. 273(5): p. 2799–807.

33. Sirisena, D.M., et al., The rfaC gene of Salmonella typhimurium. Cloning, sequencing, and enzymatic function in heptose transfer to lipopolysaccharide. Journal of Biological Chemistry, 1992. 267(26): p. 18874–18884.

34. Gronow, S., W. Brabetz, and H. Brade, Comparative functional characterization in vitro of heptosyltransferase I (WaaC) and II (WaaF) from Escherichia coli. European Journal of Biochemistry, 2000. 267(22): p. 6602–6611.

35. Czyzyk, D.J., C. Liu, and E.A. Taylor, Lipopolysaccharide biosynthesis without the lipids: recognition promiscuity of Escherichia coli heptosyltransferase I. Biochemistry, 2011. 50(49): p. 10570–2.

36. Mulichak, A.M., et al., Structure of the TDP-epi-vancosaminyltransferase GtfA from the chloroeremomycin biosynthetic pathway. Proceedings of the National Academy of Sciences, 2003. 100: p. 9238–9243.

37. Vetting, M.W., P.A. Frantom, and J.S. Blanchard, Structural and enzymatic analysis of MshA from Corynebacterium glutamicum: substrate-assisted catalysis. J Biol Chem, 2008. 283(23): p. 15834–44.

38. Sheng, F., et al., The crystal structures of the open and catalytically competent closed conformation of Escherichia coli glycogen synthase. J Biol Chem, 2009. 284(26): p. 17796–807.

39. Cote, J.M., et al., The Stories Tryptophans Tell: Exploring Protein Dynamics of Heptosyltransferase I from Escherichia coli. Biochemistry, 2017. 56(6): p. 886–895.

40. Cote, J.M., et al., Opposites Attract: Escherichia coli Heptosyltransferase I Conformational Changes Induced by Interactions between the Substrate and Positively Charged Residues. Biochemistry, 2020. 59(34): p. 3135–3147.

41. Czyzyk, D.J., et al., Escherichia coli heptosyltransferase I: investigation of protein dynamics of a GT-B structural enzyme. Biochemistry, 2013. 52(31): p. 5158–60.

42. Ramirez-Mondragon, C.A., et al., Conserved Conformational Hierarchy across Functionally Divergent Glycosyltransferases of the GT-B Structural Superfamily as Determined from Microsecond Molecular Dynamics. Int J Mol Sci, 2021. 22(9).

43. Ashkenazy, H., et al., ConSurf 2016: an improved methodology to estimate and visualize evolutionary conservation in macromolecules. Nucleic acids research, 2016. 44(W1): p. W344–W350.

44. Berezin, C., et al., ConSeq: the identification of functionally and structurally important residues in protein sequences. Bioinformatics, 2004. 20(8): p. 1322–1324.

45. Thompson, J.D., D.G. Higgins, and T.J. Gibson, CLUSTAL W: improving the sensitivity of progressive multiple sequence alignment through sequence weighting, position-specific gap penalties and weight matrix choice. Nucleic acids research, 1994. 22(22): p. 4673–4680.

46. Van Der Spoel, D., et al., GROMACS: fast, flexible, and free. J Comput Chem, 2005. 26(16): p. 1701–18.

47. Pronk, S., et al., GROMACS 4.5: a high-throughput and highly parallel open source molecular simulation toolkit. Bioinformatics, 2013. 29(7): p. 845–54.

48. Hornak, V., et al., Comparison of multiple Amber force fields and development of improved protein backbone parameters. Proteins, 2006. 65(3): p. 712–25.

49. Olsson, M.H.M., et al., PROPKA3: Consistent Treatment of Internal and Surface Residues in Empirical pKa Predictions. Journal of Chemical Theory and Computation, 2011. 7(2): p. 525–537.

50. Søndergaard, C.R., et al., Improved Treatment of Ligands and Coupling Effects in Empirical Calculation and Rationalization of pKa Values. Journal of Chemical Theory and Computation, 2011. 7(7): p. 2284–2295.

51. Jorgensen, W.L., et al., Comparison of simple potential functions for simulating liquid water. The Journal of Chemical Physics, 1983. 79(2): p. 926–935.

52. Berendsen, H.J.C., et al., Molecular dynamics with coupling to an external bath. The Journal of Chemical Physics, 1984. 81(8): p. 3684–3690.

53. Bussi, G., D. Donadio, and M. Parrinello, Canonical sampling through velocity rescaling. The Journal of Chemical Physics, 2007. 126(1): p. 014101.

54. Hess, B., et al., LINCS: A linear constraint solver for molecular simulations. Journal of Computational Chemistry, 1997. 18(12): p. 1463–1472.

55. Jakalian, A., D.B. Jack, and C.I. Bayly, Fast, efficient generation of high-quality atomic charges. AM1-BCC model: II. Parameterization and validation. Journal of Computational Chemistry, 2002. 23(16): p. 1623–1641.

56. Wang, J., et al., Development and testing of a general amber force field. Journal of Computational Chemistry, 2004. 25(9): p. 1157–1174.

57. Pearlman, D.A., et al., AMBER, a package of computer programs for applying molecular mechanics, normal mode analysis, molecular dynamics and free energy calculations to simulate the structural and energetic properties of molecules. Computer Physics Communications, 1995. 91(1): p. 1–41.

58. Sousa da Silva, A.W. and W.F. Vranken, ACPYPE - AnteChamber PYthon Parser interfacE. BMC research notes, 2012. 5: p. 367–367.

59. Grant, B.J., L. Skjærven, and X.-Q. Yao, The Bio3D packages for structural bioinformatics. Protein Science, 2021. 30(1): p. 20–30.

60. David, C.C. and D.J. Jacobs, Principal Component Analysis: A Method for Determining the Essential Dynamics of Proteins, in Protein Dynamics: Methods and Protocols, D.R. Livesay, Editor. 2014, Humana Press: Totowa, NJ. p. 193–226.

61. Eargle, J. and Z. Luthey-Schulten, NetworkView: 3D display and analysis of protein:RNA interaction networks. Bioinformatics (Oxford, England), 2012. 28(22): p. 3000–3001.

62. Scarabelli, G. and B.J. Grant, Mapping the structural and dynamical features of kinesin motor domains. PLoS computational biology, 2013. 9(11): p. e1003329–e1003329.

63. Kollman, P.A., et al., Calculating Structures and Free Energies of Complex Molecules: Combining Molecular Mechanics and Continuum Models. Accounts of Chemical Research, 2000. 33(12): p. 889–897.

64. Wang, E., et al., End-Point Binding Free Energy Calculation with MM/PBSA and MM/GBSA: Strategies and Applications in Drug Design. Chemical Reviews, 2019. 119(16): p. 9478–9508.

65. Wang, C., et al., Recent Developments and Applications of the MMPBSA Method. Frontiers in molecular biosciences, 2018. 4: p. 87–87.

66. Genheden, S. and U. Ryde, The MM/PBSA and MM/GBSA methods to estimate ligand-binding affinities. Expert opinion on drug discovery, 2015. 10(5): p. 449–461.

67. Miller, B.R., 3rd, et al., MMPBSA.py: An Efficient Program for End-State Free Energy Calculations. J Chem Theory Comput, 2012. 8(9): p. 3314–21.

68. Harms, M.J., et al., The pKa Values of Acidic and Basic Residues Buried at the Same Internal Location in a Protein Are Governed by Different Factors. Journal of Molecular Biology, 2009. 389(1): p. 34–47.

69. Lemieux, R.U., How Water Provides the Impetus for Molecular Recognition in Aqueous Solution. Accounts of Chemical Research, 1996. 29(8): p. 373–380.

70. Hayward, S. and H.J.C. Berendsen, Systematic analysis of domain motions in proteins from conformational change: New results on citrate synthase and T4 lysozyme. Proteins: Structure, Function, and Bioinformatics, 1998. 30(2): p. 144–154.

71. Hayward, S. and R.A. Lee, Improvements in the analysis of domain motions in proteins from conformational change: DynDom version 1.50. Journal of Molecular Graphics and Modelling, 2002.21(3): p. 181–183.

72. Nkosana, N.K., et al., Synthesis, kinetics and inhibition of Escherichia coli Heptosyltransferase I by monosaccharide analogues of Lipid A. Bioorganic & medicinal chemistry letters, 2018. 28(4): p. 594–600.

