## Supplemental Information for "Ligand Induced Conformational and Dynamical Changes in a GT-B Glycosyltransferase: Molecular Dynamic Simulations of Heptosyltransferase I Apo, Binary and Ternary Complexes"

| Simulation | PDB Code | ADP-Hep | FDLA | ADP | FDHLA |
|---|---|---|---|---|---|
| Apo | 2GT1 | N/A | | | |
| ADP-Hep•FDLA•HepI (Substrates) | 6DFE | Added | Present | N/A | |
| FDLA•HepI (Substrate) | 6DFE | N/A | Present | | |
| ADP-Hep•HepI (Substrate) | 2H1H | Modified | N/A | | |
| ADP•FDHLA•HepI (Products) | 6DFE | N/A | | Added | Modified |
| FDHLA•HepI (Products) | 6DFE | | | N/A | Modified |
| ADP•HepI (Products) | 2H1F | | | Present | N/A |

**Supplemental Table 1:** The crystal structures used for simulations and details on whether a ligand was present in the original structure or if it was added. The fluorinated ADP-Hep was taken from PDB:2H1H and the fluorine was replaced with a hydroxyl group in the proper stereoconfiguration. The FDHLA was constructed by the addition of a Heptose on the FDLA in the right stereo/regioconfiguration.

| Simulation | RMSD (Å) | $R_{gyr}$ (Å) | Interdomain Center of Mass Distance (Å) | RMSF (Residues >1.5 Å) |
|---|---|---|---|---|
| Apo | 1.70 ± 0.25 | 21.11 ± 0.16 | 29.70 ± 0.46 | N3 (61-68), C2 (216-219,230), C6(316-320) |
| ADP-Hep•HepI (Substrate) | 1.82 ± 0.29 | 21.04 ± 0.15 | 29.34 ± 0.42 | N3 (58-71), C1 (206), C2 (230), C5 (279-280), C6 (299,316,318-320) |
| FDLA•HepI (Substrate) | 1.82 ± 0.22 | 20.99 ± 0.11 | 29.37 ± 0.34 | N3 (58,61-68), C6 (319,320) |
| ADP-Hep•FDLA•HepI (Substrates) | 1.75 ± 0.31 | 21.21 ± 0.12 | 30.01 ± 0.37 | N3 (62-67), C1 (188-189), C5 (283-284) C6 (299, 317-320) |
| ADP•FDHLA•HepI (Products) | 1.85 ± 0.31 | 21.36 ± 0.23 | 28.62 ± 0.54 | N3 (61-68), N4 (103), N6 (135,156), C1 (188-189,206), C2 (218), C5 (280-281,287-291), C6 (298-318) |
| ADP•HepI (Products) | 1.63 ± 0.29 | 21.08 ± 0.13 | 29.45 ± 0.37 | N3 (58, 59, 61-68), C1 (206), C2 (230), C5 (284, 287), C6 (298-299,316-318) |
| FDHLA•HepI (Products) | 1.87 ± 0.28 | 21.10 ± 0.14 | 28.53 ± 0.42 | N3 (61-68), C1 (206), C2 (218-219,230)<br>C5 (284,287), C6 (299,315-318) |
| ADP•FDHLA•HepI (Products) (D13+H) | 2.20 ± 0.28 | 21.02 ± 0.17 | 28.20 ± 0.39 | N3 (61-71), C1 (188-190,206) C2 (217-219,230), C3 (239), C5 (279, 280), C6 (296-301, 316-318) |

**Supplemental Table 2:** Average values of RMSD and radius of gyration from the length of one representative simulation with standard deviations. RMSF values are reported for regions and residues that have a greater than 1.5 Å fluctuation.

| Residue | Apo | ADP-Hep•HepI | FDLA•HepI | ADP-Hep•FDLA•HepI | ADP•FDHLA•HepI | ADP•HepI | FDHLA•HepI |
|---|---|---|---|---|---|---|---|
| Lys7* | 9.27 | 9.54 | 8.86 | 8.81 | 5.77 | 9.42 | 5.78 |
| Asp13* | 4.46 | 4.48 | 6.72 | 6.79 | 10.38 | 4.40 | 10.36 |
| Glu38 | 6.06 | 6.23 | 6.49 | 6.72 | 6.86 | 6.22 | 6.69 |
| His168 | 5.93 | 6.38 | 6.38 | 6.38 | 5.93 | 5.93 | 5.93 |
| His198 | 6.68 | 6.63 | 6.63 | 6.63 | 6.68 | 6.68 | 6.68 |
| Glu222 | 3.47 | 5.15 | 4.10 | 5.19 | 4.48 | 4.40 | 3.52 |
| Asp261* | 4.11 | 6.01 | 5.58 | 6.28 | 5.81 | 5.20 | 5.45 |
| His266* | 4.76 | 3.67 | 3.24 | 3.47 | 2.78 | 3.66 | 3.13 |

**Supplemental Table 3:** pKa of ionizable sidechains that have a greater than 0.5 pKa shift relative to apo or within 0.5 pH units of 7.0 as determined by PROPKA3.

| PCA Covar. | Apo | ADP-Hep•FDLA•HepI | ADP-Hep•HepI | FDLA•HepI | ADP•FDHLA•HepI | ADP•HepI | FDHLA•HepI | ADP•FDHLA•HepI (Products) (D13+H) |
|---|---|---|---|---|---|---|---|---|
| PC1 (%) | 19.2 | 30.2 | 26.4 | 20 | 29.9 | 31.4 | 17.1 | 40.7 |
| PC2 (%) | 13.9 | 11.5 | 14.7 | 10.5 | 14.1 | 13.1 | 15.8 | 11.6 |
| PC3 (%) | 11.1 | 5.7 | 9.7 | 8.5 | 10.7 | 7 | 9.3 | 5 |
| Total (%) | 44.6 | 47.4 | 50.8 | 39 | 54.7 | 51.5 | 42.2 | 62.3 |

**Supplemental Table 4:** PCA percent covariance table

| Path | Apo | ADP-Hep•FDLA• HepI | ADP-Hep• HepI | FDLA• HepI | ADP•FDHLA• HepI | ADP• HepI | FDHLA• HepI | ADP•FDHLA• HepI (Products) (D13+H) |
|---|---|---|---|---|---|---|---|---|
| 60->120 | N3(58, 60, 61, 62); N5(120); C1(195, 196, 197); C2(224, 225, 226, 228) | N3(59, 60); N5(119, 120, 121); C1(190, 191, 192, 193, 194, 195) | N1(4); N2(34, 35, 36, 37, 38); N3(56, 57, 58, 59, 60, 61); N4(90); N5(111, 112, 113, 114, 119, 120) | N1(7, 8); N2(36, 37, 38); N3(57, 58, 59, 60, 61); N4(94, 95, 96); N5(120) | N1(8, 9); N2(37, 38); N3(58, 59, 60, 61); N4(96, 97); N5(120, 121); C1(185) | N1(7 , 8); N2(38); N3(58, 59, 60, 61); N4(94, 95, 96); N5(120, 121); C1(193, 194, 195) | N1(5); N2(36, 37); N3(57, 58, 59, 60); N4(91, 92); N5(112, 113, 114, 119, 120) | N3(60); N5(120, 121); C1(194, 195, 196, 197, 198, 199, 200) |
| 60->192 | N3(58, 60, 61, 62); C1 ( 192, 193, 194, 195, 196); C2 (225, 226, 228) | N3(59, 60 61, 62, 63); C1(189, 190, 191, 192, 193, 194, 195) | N3(59, 60, 61, 62, 63, 64); C1(190, 191, 192, 193) | N3(58, 60); C1(192, 193, 194, 195, 196); C2(223, 224, 225, 226, 228) | N3(58, 60, 61); C1(185, 186, 187, 191, 192, 193, 194, 195, 196); C2(224, 225, 226, 228) | N2(38); N3(58, 59, 60, 61); C1(192, 193, 194, 195, 196, 197) | N3(57, 58, 59, 60); C1(183, 184, 185, 187, 189, 192, 193); C2(215); C4(260) | N3(60) C1(192, 193, 194, 195, 196, 197, 198, 199) |
| 60->242 | N3(58, 59, 60, 61, 62, 64, 65, 68, 71, 72, 73, 74); C3(238, 239, 240, 241, 242) | N3(58, 60, 61); C2(223, 224, 226, 227); C3(238, 239, 240, 241, 242) | N3(58, 59, 60, 61, 62); C2(223, 224); C3(239, 240, 241, 242) | N3(58, 60); C2(223, 226, 227) C3(238, 239, 240, 241, 242) | N3(58, 59, 60, 61, 62); C3(238, 239, 240, 241, 242) | N3(59, 60, 61, 62, 70, 71, 72, 73, 74, 75); C2(223) C3(240, 241, 242) | N3(56, 57, 58, 59, 60, 61); C2(215); C3(238, 239, 240, 241, 242 | N3(60); C1(200, 201, 202); C3(242, 243, 244, 245, 246) |
| 60->261 | N3(60, 61, 62); C1(185, 186, 187, 194, 195, 196); C2(225, 226, 228); C4(260, 261) | N3(59, 60); C1(190, 191, 192, 193, 194, 195); C4(260, 261); C5(279, 280, 281) | N3(59 , 60, 61, 62, 63); C1(187, 189, 193, 194); C4(260, 261) | N3(58, 60); C1(193, 194, 195, 196); C2(223, 224, 225, 226, 227, 228); C4(260 261) | N2(38, 39, 40, 41); N3(58, 60, 61); C4(261, 262, 263, 264, 266, 267) | N2(38); N3(58, 59, 60, 61); C1(192, 193, 194, 195, 196); C4(260, 261) | N3(56, 57, 58, 59, 60, 61); C1(183, 184); C2(215, 216); C3(238); C4(259, 260, 261) | N3(60); C1(193, 194, 195, 196, 197, 198, 199); C4(260, 261) |
| 120->192 | N4(95, 96, 97); N5(120, 121, 124); C1(192, 193, 194, 195) | N5(118, 119, 120, 121); C1(191, 192); C5(279, 280, 281, 282) | N1(4); N2(34, 35, 36, 37, 38); N3(57, 58, 59, 60, 61, 62); N4(90); N5(112, 113, 114, 119, 120); C1(191, 192, 193) | N4(96, 97, 100, 101, 105, 106, 107); N5(120, 121, 122, 124); C1(191, 192, 193); C5(281) | N4(96); N5(119, 120, 121, 122, 123, 124); C1(191, 192, 193) | N5(116, 118, 119, 120, 121, 122, 123, 124); C1(192) | N5(114, 115, 116, 118, 119, 120, 121); C1(192, 193); C4(260); C5(278) | N5(116, 119, 120, 121, 122, 123, 124, 125); C1(192); |
| 120->242 | N4(96); N5(120, 121); C1(195, 196, 197); C2(222, 223, 224, 225, 226); C3(238, 239, 240, 241, 242) | N5(114, 115, 119, 120); N6(132, 133); C3(237, 238, 239, 240, 241, 242) | N1(15, 16, 18); N2(47); N5(114, 115, 118, 119, 120); N6(133, 134, 141, 142, 143, 165); C3(242, 243, 245) | N1(7, 8, 9, 10, 11); N2(37, 38); N3(57); N4(94); N5(120); C2(223, 226, 227); C3(238, 239, 240, 241, 242, 248); C4(267, 271) | N4(96, 97, 98, 100, 101); N5(120, 121, 122, 123, 124); C3(242, 243, 244) | N5(120); C1(186, 193, 194, 195, 196); C2(216, 223, 224, 225, 226, 227, 228); C3(238, 239, 240, 241, 242) | N5(114, 119, 120); C1(184); C2(215); C3(238, 239, 240, 241, 242); C4(259, 260) | N5(120); C3(242, 243, 244, 245, 246, 247, 248, 249); C6(308, 309) |
| 120->261 | N4(96); N5(120, 121, 122, 123, 124, 125); C4(260, 261) | N5(114, 115, 116, 117, 118, 119, 120, 121); N6(132); C4(261); C5(279) | N1(15); N5(114, 115, 118, 119, 120); N6(133, 134, 141, 142, 143); C4(261, 262, 263, 265, 266, 267) | N1(7, 9, 10, 11, 12, 14); N4(94, 95); N5(120); C4(261, 262, 263, 264, 265, 266, 267) | N4(96, 97, 100, 104); N5(119, 120, 121, 122, 123, 124); C4(259, 260, 261); | N5(118, 119, ,120, 121, 122); C1(192, 193, 194); C4(260, 261); C5(278) | N5(113, 114, 115, 116, 118, 119, 120, 121); C4(260, 261); | N5(119, 120, 121, 122, 123, 124); C4(260, 261) |
| 13->60 | N1(13) N3(58, 60, 61); C1(196, 197, 198, 199); C2(225, 226, 228) | N1(13, 14, 16, 17, 18, 19, 20, 21, 23, 24); N3(58, 60, 61); N6(149) | N1(13 , 14, 15); N3(59, 60, 61, 62, 63); C1(187, 188, 189, 190); C4(266, 267) | N1(7, 8, 13); N2(36, 37, 38); N3(56, 57, 58, 59, 60, 61) | N1(8, 9, 10, 12, 13, 14); N2(37, 38, 41); N3(58, 59, 60, 61) | N1(8 , 9, 10, 11, 12, 13); N2(38, 39); N3(57, 58, 59, 60, 61) | N1(8, 9, 10, 13, 14); N2(37, 38); N3(56, 57, 58, 59, 60, 61) | N1(8, 10, 13); N3(58, 59, 60, 63); C1(197, 198, 199, 200, 201, 202) |
| 13->120 | N1(12, 13); N5(120, 121); C1(195, 196, 197, 198, 199, 200) | N1(13, 16, 17, 20); N5(114, 115, 118, 119, 120); N6(132, 133 146, 147) | N1(13, 16); N5(114, 115, 118, 119, 120); N6(133, 134, 140, 141, 142, 143) | N1(7, 8, 9, 10, 11, 12, 13); N4(94, 95, 96); N5(119, 120, 121) | N1(9, 10, 11, 12, 13, 14); N4(95, 96); N5(119, 120, 121) | 13, 94, 95, 96, 118, 119, 120, 121, 122, 123, 124 | 13, 16, 114, 119, 120, 132, 133, 141, 142, 143, 144, 145, 146, 147 | 13, 93, 94, 95, 96, 118, 119, 120, 121, 122 |
| 13->192 | N1(13); C1(192, 193, 194, 195, 196, 197, 198, 199) | N1(13, 16, 17, 20); N6(146, 147); C1(192); C5(279, 280, 281, 282, 283); C6(298) | 13 , 14, 15, 187, 188, 189, 190, 191, 192, 193, 266, 267 | 12, 13, 14, 15, 192, 193, 260, 261, 262, 263, 264, 265, 266, 267 | N1(9, 10, 11, 12, 13); C1(191, 192, 193, 194, 195, 196) | N1(13); C1(192 193, 194, 195, 196, 197, 198, 199, 200, 201); | N1(12, 13, 14, 15); C1(187, 192); C4(261, 262, 263, 265, 266, 267) | N1(13); N4(94, 95, 96); N5(121, 122, 123, 124, 125); C1(192); |
| 13->242 | N1(12, 13); C1(196, 197, 198, 199); C2(223, 224, 225) C3(241, 242, 243, 244, 245, 248); C4(267, 271) | N1(13, 16, 17, 18, 19, 20) N3(74); N6(146, 148, 149, 150, 163); C3(237, 238, 239, 240, 241, 242, 243, 244, 245, 246, 248); C4(270, 271) | N1(13, 14, 15, 17, 18); N2(47, 48); N6(165); C3(242, 243, 244, 245, 246); C4(267, 271) | N1(12, 13, 14, 15); C3(242, 243, 244, 245, 246, 247, 248); C4(267, 268, 270, 271) | N1(12, 13, 14, 15, 16); C3(242, 243, 244, 245); C4(267, 270) | N1(12, 13); C1(196, 197, 200, 201); C2(223, 224, 225, 227, 228, 229); C3(238, 239, 240, 241, 242, 243, 244, 245, 246, 248); C4(271) | N1(12, 13, 14, 15); C3(242, 243, 244, 245, 246, 248); C4(267, 270, 271) | N1(10, 11, 12, 13); C3(241, 242, 243, 244, 245, 246) |
| 13->261 | N1(13); C1(193, 194, 195, 196, 197, 198, 199); C4(260, 261) | N1(13, 16, 17, 18, 20); N6(132, 133, 146, 147, 150); C4(261) | N1(13 , 14, 15, 16); C4(260, 261, 262, 263, 264, 265, 266, 267) | N1(12, 13, 14, 15); C4(261, 262, 263, 264, 265, 266, 267) | N1(12, 13, 14, 15, 16) C4(261, 262, 263, 264, 266, 267) | N1(13); C1(193, 194, 195, 196, 197, 198, 199, 200, 201); C4(260, 261) | N1(12, 13, 14, 15); C4(261, 262, 263, 265, 266, 267) | N1(10, 11, 12, 13, 14, 15); C4(261, 262, 263, 264, 266) |

**Supplemental Table 5:** Consensus residues involved in each of the collection of shortest paths of communication in their respective simulations.

A)

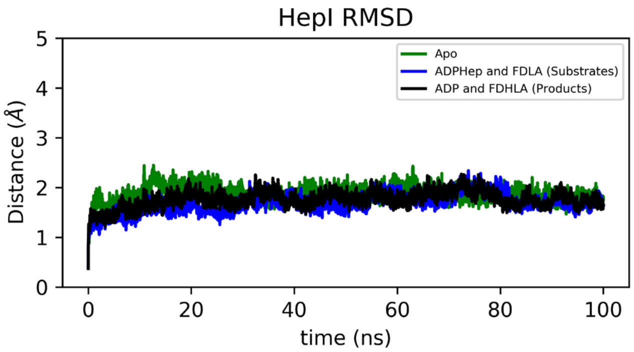


B)

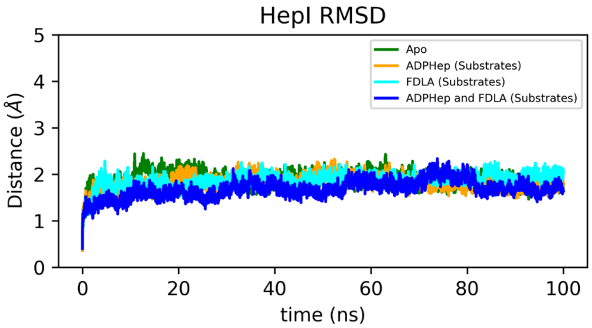


C)

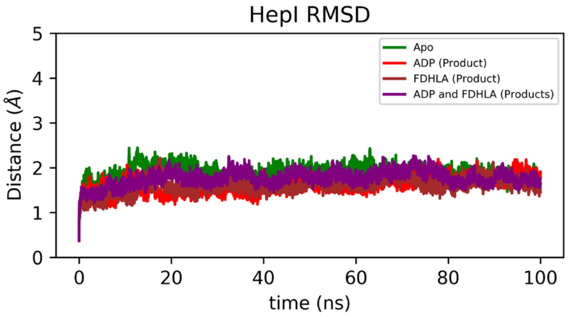


**Supplemental Figure 1:** (**A**) Backbone RMSD of HepI apo, substrate ternary complex and product ternary complex with deprotonated Asp13. (**B**) Backbone RMSD of HepI apo, substrate binary complexes (ADP-Hep•HepI, FDLA•HepI) and substrate ternary complex. (**C**) Backbone RMSD of HepI apo, product binary complexes (ADP•HepI, FDHLA•HepI) and product ternary complex with deprotonated Asp13.

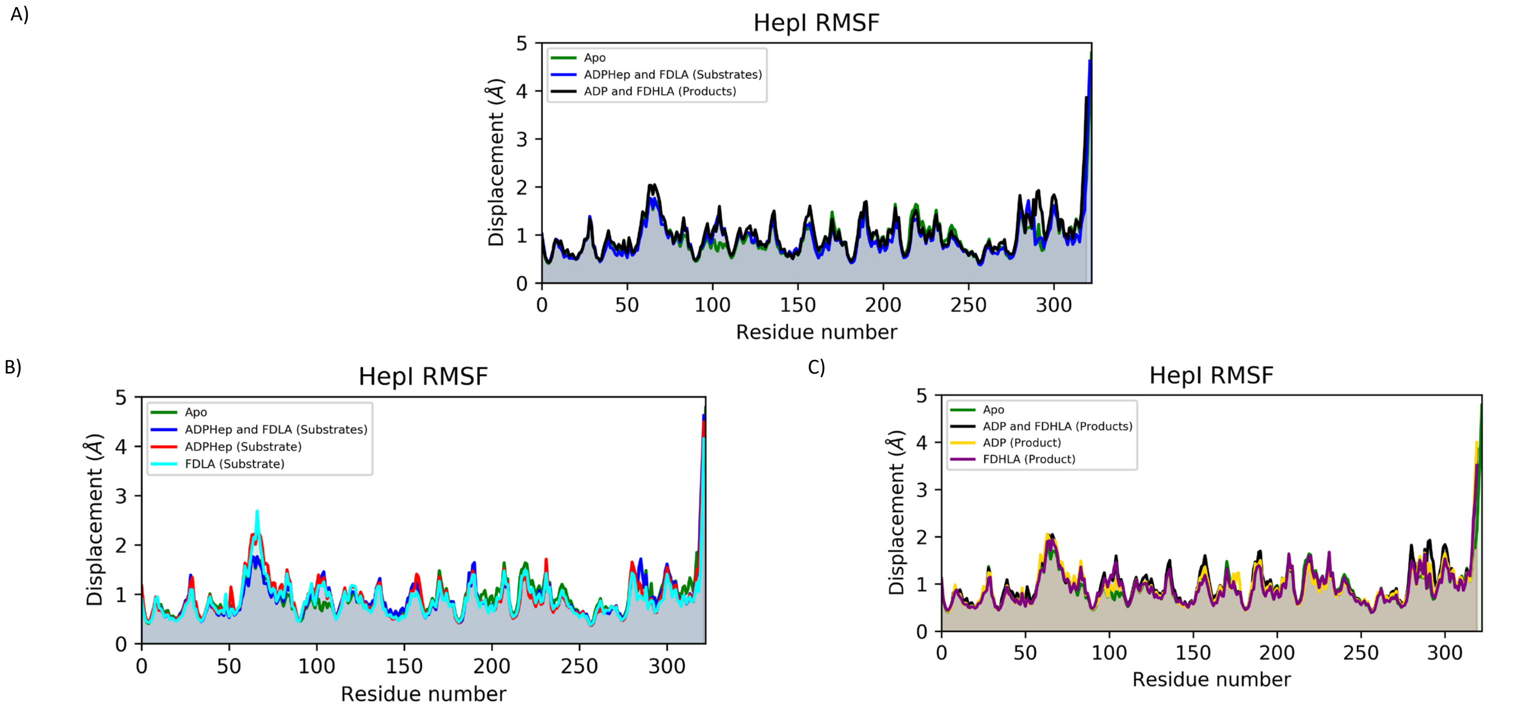


**Supplemental Figure 2:** (**A**) Cα RMSF of HepI apo, substrate ternary complex and product ternary complex with deprotonated Asp13. (**B**) Cα RMSF of HepI apo, substrate ternary complex and substrate binary complexes (ADP-Hep•HepI, FDLA•HepI). (**C**) Cα RMSF of HepI apo, product ternary complex with deprotonated Asp13 and product binary complexes (ADP•HepI, FDHLA•HepI).

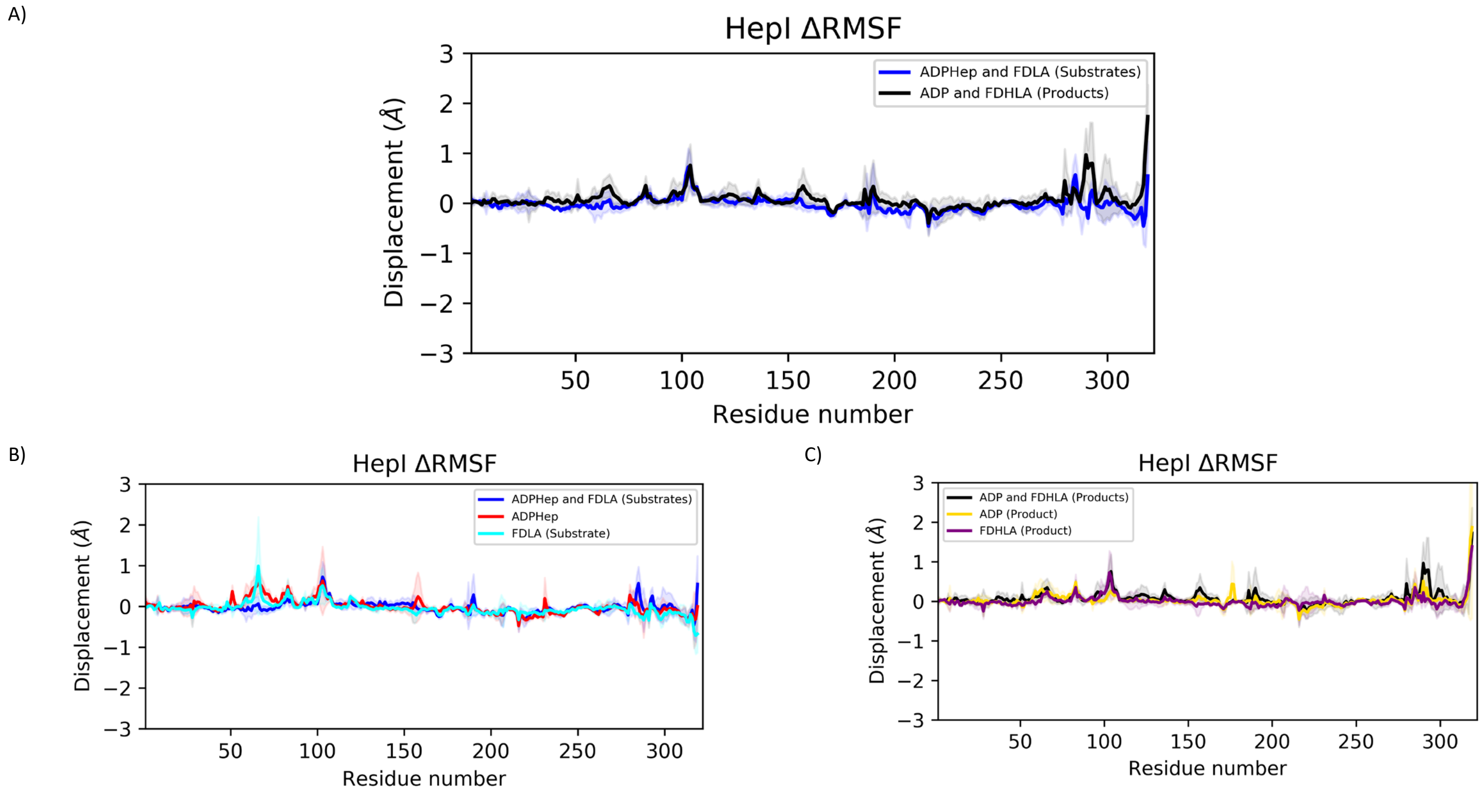


**Supplemental Figure 3:** (**A**) Cα ΔRMSF of HepI substrate ternary complex and product ternary complex with deprotonated Asp13. (**B**) Cα ΔRMSF of HepI substrate ternary complex and substrate binary complexes (ADP-Hep•HepI, FDLA•HepI). (**C**) Cα ΔRMSF of HepI product ternary complex with deprotonated Asp13 and product binary complexes (ADP•HepI, FDHLA•HepI). solid lines are average differences relative to apo (i.e. $RMSF_{substrates}$-$RMSF_{apo}$) and shaded region is the standard deviation of the average difference. Positive values indicate those residues are more flexible relative to HepI apo.

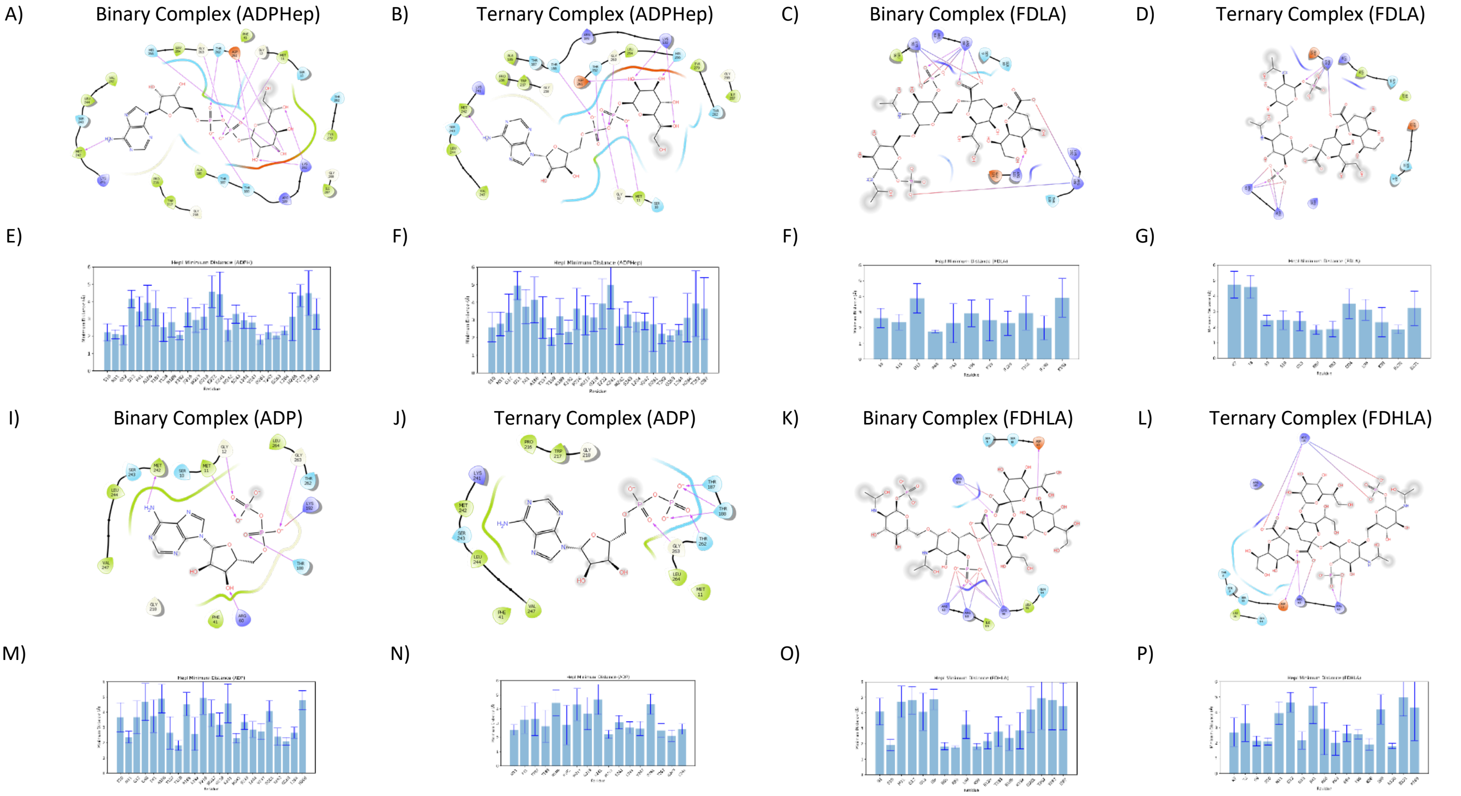


**Supplemental Figure 4:** Ligand interaction diagram of (**A-B**) ADP-Hep, (**B-C**) FDLA, (**I-J**) ADP and (**K-L**) FDHLA from binary and ternary complexes. Bar plots of residues with average minimum distances of less than 5 Å from (**E-F**) ADP-Hep, (**G-H**) FDLA, (**M-O**) ADP and (**P-Q**) FDHLA from binary and ternary complexes.

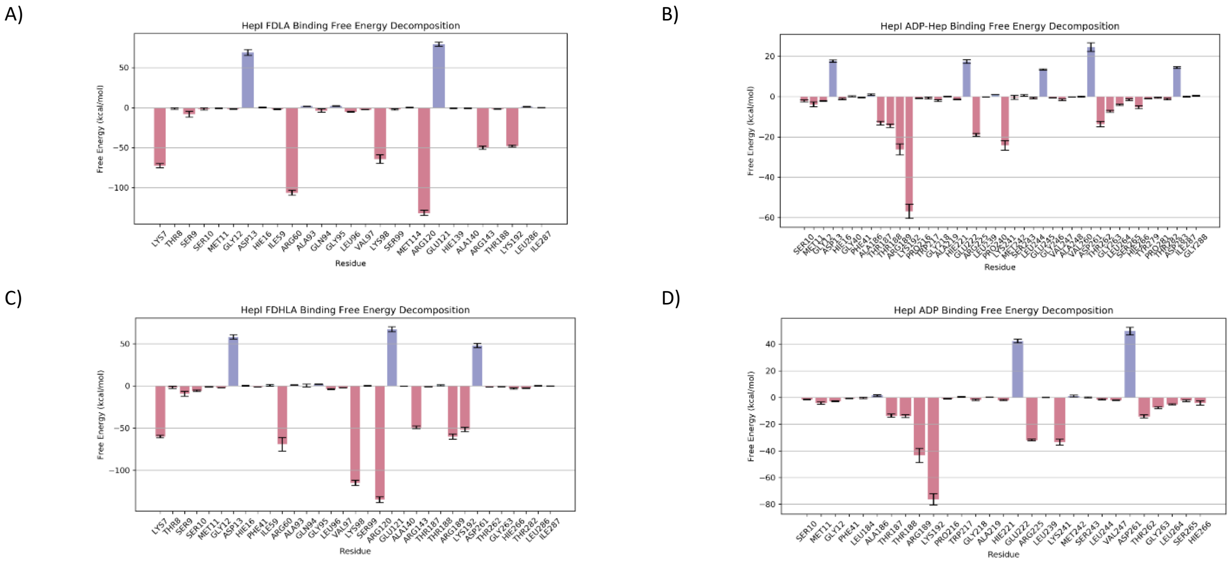


**Supplemental Figure 5:** Binding free energy contribution of residues within 6 Å of (**A**) FDLA, (**B**) ADP-Hep, (**C**) FDHLA and (**D**) ADP in the ternary complex.

A)

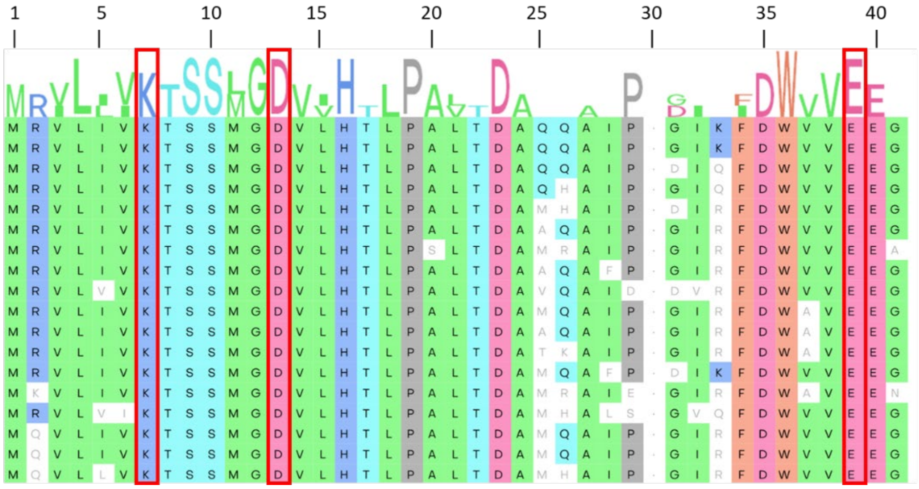


B)

| Residue | Frequency |
|---|---|
| Lys7 | 84% |
| Asp13 | 100% |
| Glu38 | 98% |
| His168 | 23% |
| His198 | 27% |
| Glu222 | 98% |
| Asp261 | 100% |
| His266 | 100% |

**Supplemental Figure 6:** (**A**) 18 representative sequences of Heptosyltransferase I from a multiple sequence alignment of 150 sequences from various homologues with residues highlighted in red boxes that demonstrate a large pKa shift. (**B**) Table of residues identified with pKa shifts and their corresponding frequency of occurrence at their respective positions in the multiple sequence alignment.

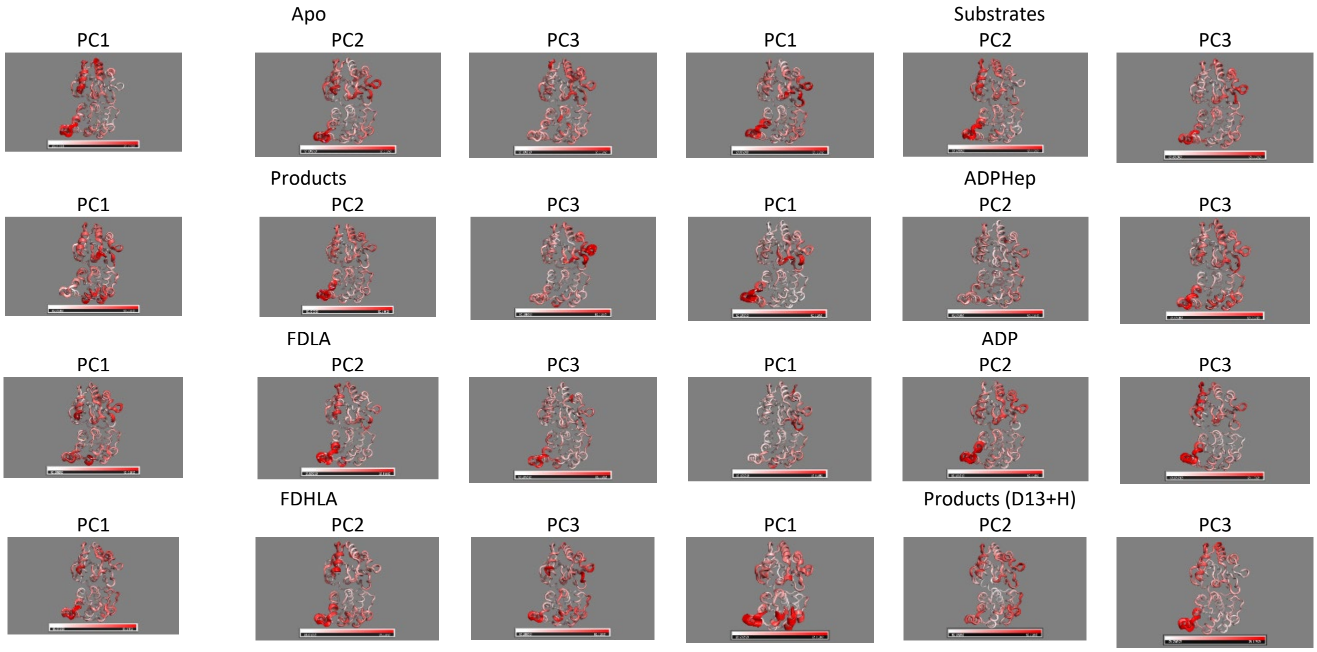


**Supplemental Figure 7:** First three principle components of (**A**) HepI apo and (**B**) ) substrate complex (**C**) ) product complex (**D**) ADP-Hep binary complex (**E**) FDLA binary complex (**F**) ADP binary complex (**G**) FDHLA binary complex and (**H**) product ternary complex with protonated Asp13.

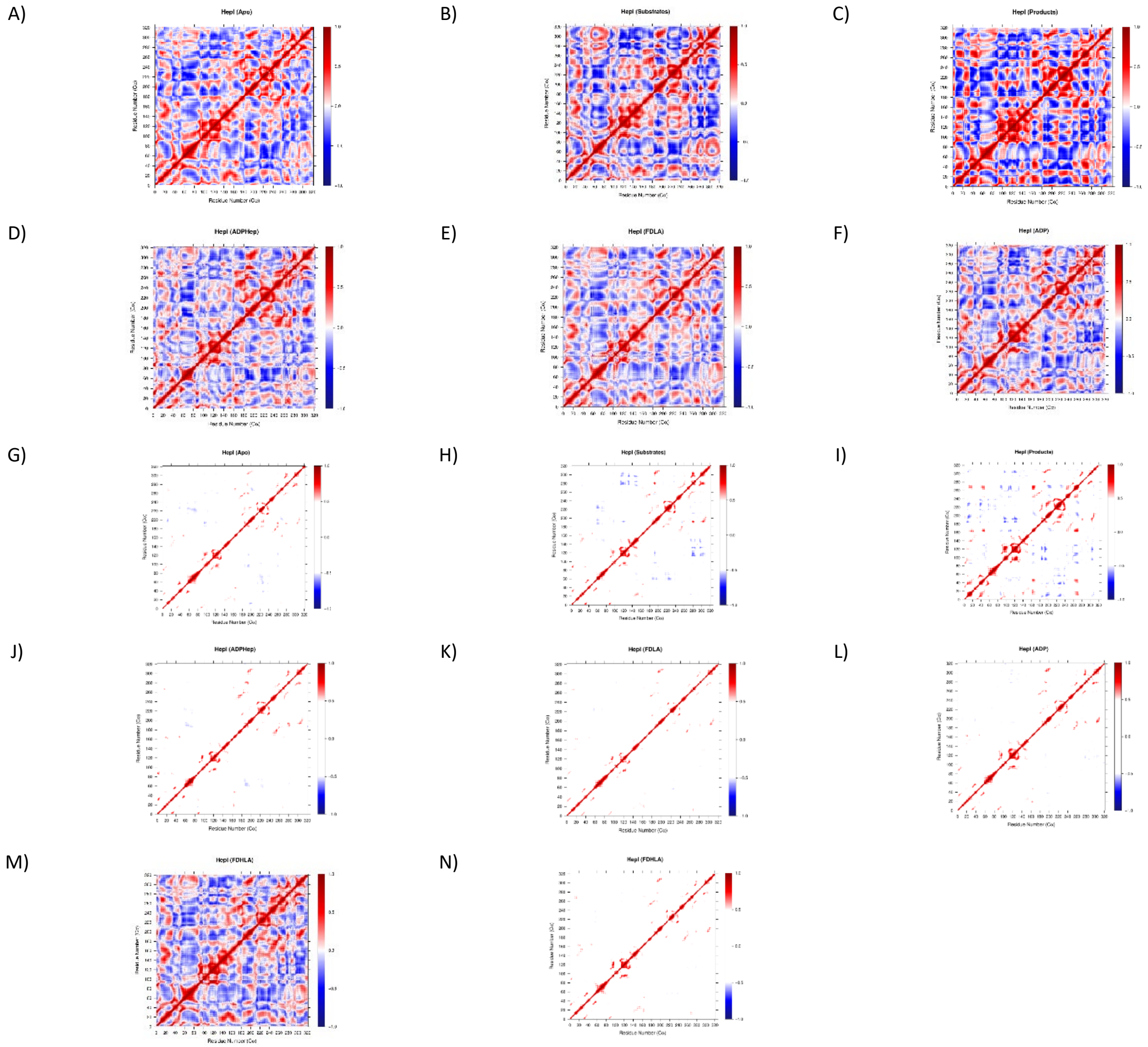


**Supplemental Figure 8:** Dynamic cross correlation map of (**A,G**) HepI apo, (**B,H**) ADP-Hep•FDLA•HepI, (**C,I**) ADP•FDHLA•HepI, (**D,J**) ADP-Hep•HepI , (**E,K**) FDLA•HepI, (**F,L**) ADP•HepI, and (**M,N**) FDHLA•HepI.

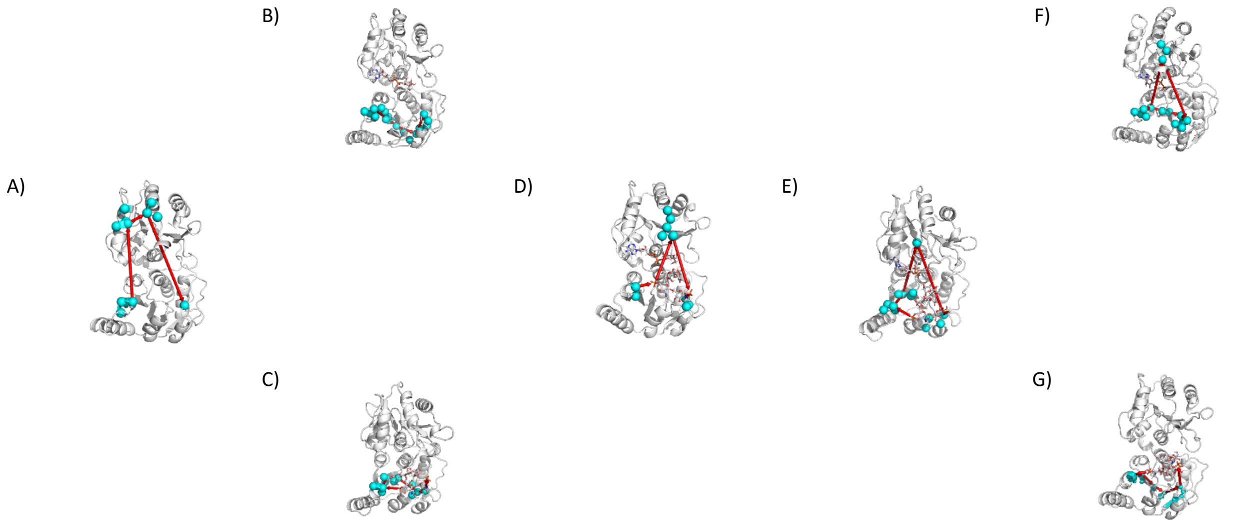


**Supplemental Figure 9:** Protein communication network between residues Arg60 and Arg120 for (**A**) HepI apo, (**B**) with ADP-Hep, (**C**) with FDLA, (**D**) with both substrates, (**E**) with both products and deprotonated Asp13, (**F**) with ADP and (**G**) with FDHLA.
